# Computational Design and Evaluation of De Novo Protein Minibinders Targeting Specific Sites of STAT3

**DOI:** 10.64898/2026.10.24.761574

**Authors:** Emily R. Sragovicz, Alexander V. Tobias

**Affiliations:** Country Day School, Alajuela Province, Costa Rica; Throop Research Academy, Gaithersburg, MD, USA

## Abstract

STAT3 is a transcription factor involved in the progression of many types of cancer, which makes it an attractive therapeutic target. Small-molecule STAT3 inhibitors have experienced a high failure rate in clinical trials, indicating alternative approaches may be warranted. To address these limitations, we used BindCraft, a modern AI-driven protein design tool, to create miniprotein binder candidates against several sites of STAT3. We selected seven target hotspots on the SH2, linker, DNA-binding and coiled-coil domains of STAT3. Of 139 designs that passed BindCraft’s filters, 11 high-confidence binders emerged from five of the seven hotspots. These candidate binders were evaluated with AlphaFold3 (AF3). All eleven are predicted to dock with their intended domain; most within 4.5 Å of one or more of their targeted hotspot residues. Several minibinders are predicted to dock simultaneously to STAT3 without overlap. Such pairs have the potential to constitute STAT3 sandwich assays. BLASTp searches found no matches with known protein sequences for six of the binders and only very low-significance matches for the other five. We disclose herein the sequences of our top binders targeting diverse regions of the STAT3 surface with the hope that experimental validation by the broader research community will yield promising leads for preclinical drug development against this important cellular regulator.

## 1. Introduction

STAT3, a member of the Signal Transducer and Activator of Transcription (STAT) family, is a eukaryotic transcription factor involved in homeostasis and has been implicated in the pathogeny of numerous diseases. STAT3 is particularly known for its role in various types of cancer [1]. Molecules such as cytokines, growth factors and oncogenes interact with STAT3. In response to cytokine signals received by cell surface receptors, STAT3 becomes activated by tyrosine phosphorylation. Activated STAT3 then travels to the nucleus, where it serves as a transcription factor by binding to specific DNA sequences. Normal cellular functions must be altered for cancer to grow and spread. Such changes include cell proliferation and angiogenesis, both of which involve STAT3 [2]. An irregular concentration of STAT3 has been reported in 70% of cancers [3]. Several human cancers are considered to have a poor prognosis when STAT3 becomes activated. These include head and neck tumors, B-cell lymphoma, cervical cancer, gastric carcinoma and colon cancer [2]. Because of its key role in cancer progression, STAT3 is an attractive therapeutic target. While the SH2 domain of STAT3 has been commonly targeted for inhibition by candidate therapeutic molecules, this domain is highly conserved among other STAT proteins, and many such ligands have lacked high selectivity for STAT3-SH2 [1]. Engineering molecules to inhibit the DNA-binding domain of STAT3 also presents challenges, as it is a large, flat surface and interfering with it may negatively affect the transcription of essential genes [1]. Overall, STAT3 inhibitors have experienced a high failure rate in clinical trials [4].

To address these limitations, we used BindCraft, a modern AI-driven protein generation tool to design minibinder protein candidates against the SH2 domain, linker domain, DNA-binding domain and coiled-coil domain of STAT3. We hypothesized that BindCraft could help us design de novo miniproteins that bind to STAT3 at or near the input hotspot amino acids (according to AlphaFold evaluations [5]), and that their sequences would be completely novel, with no significant similarity to any other known proteins, as assessed by BLAST searches. From this work, we generated 11 promising sequences for protein minibinders to diverse portions of the STAT3 surface. They are disclosed herein so that others may validate them experimentally, thereby providing new leads for therapeutics that target this important cellular regulator.

## 2. Materials and Methods

### 2.1 Experimental and Technical Design

Candidate STAT3 binders were generated using the BindCraft [6] pipeline running in Google Colaboratory. Suitable “hotspot” regions of the STAT3 surface for binder targeting were found with the assistance of ChimeraX protein visualization software. Each selected hotspot was then input into the BindCraft pipeline for binder generation. BindCraft “campaigns” were run 1–4 times per hotspot. The number of campaigns depended on how favorable the hotspot appeared as a target (how accessible it was) and on the success of the initial campaign(s), with less success prompting additional campaigns. Binders with interface predicted template modeling (ipTM) scores of ≥ 0.60 from the BindCraft pipeline (which runs AlphaFold2 (AF2)) were further evaluated with AF3 on the public AlphaFold Server (AFS). Minbinder protein sequences with AF3 ipTM scores ≥ 0.75 and pLDDT values ≥ 90 for ≥ 85% of the protein-binder complex were considered high-confidence. High-confidence binders were visualized with ChimeraX software to determine which portions of STAT3 interact with and are covered by the binder, especially in relation to the intended hotspot. Finally, BLASTp searches were conducted to assess sequence novelty. Figure 1 depicts the workflow followed in this study.

**Figure 1.**
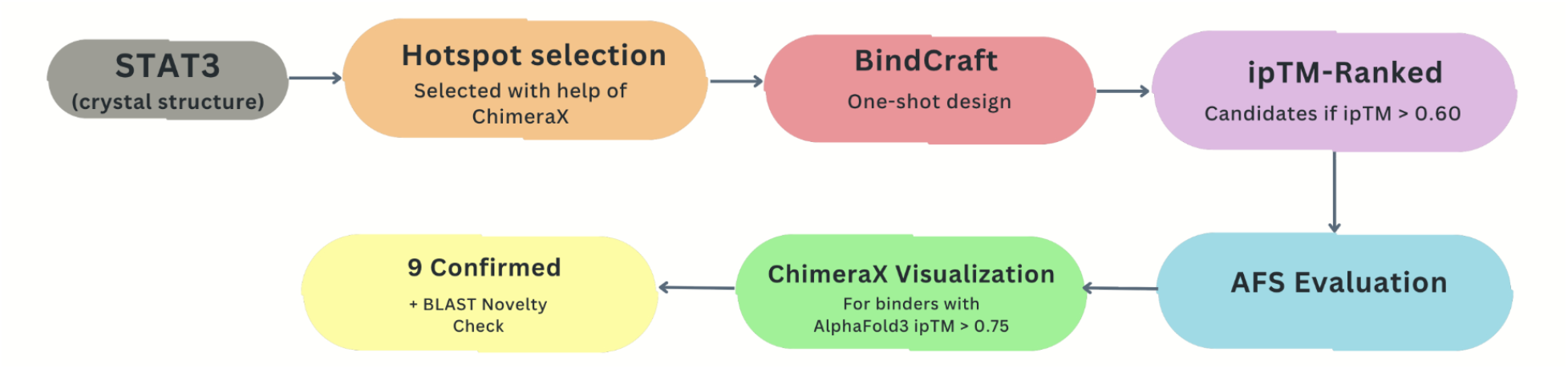
Workflow used in this study.

**Figure 2.**
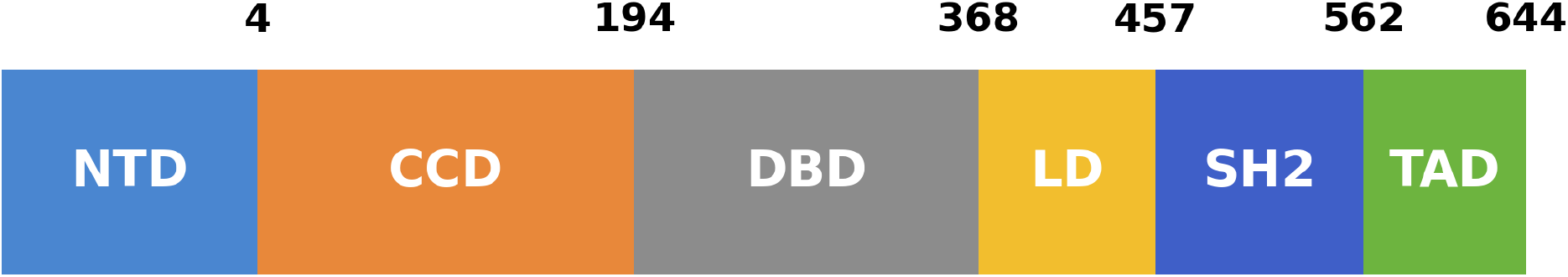
Domains of STAT3 referenced in Tables 5 and 6. NTD, N-terminal domain; CCD, coiled-coil domain; DBD, DNA-binding domain; LD, linker domain; SH2, Src homology 2 domain; TAD, transactivation domain. Numbers refer to the approximate amino acids (AFS numbering) at domain borders.

### 2.2 Target structure and hotspot selection

Hotspots were chosen based on five criteria. First, each hotspot was limited to 3 amino acids (3 as the preferred number but 2 still acceptable) to balance identification of a specific region without overspecifying, to allow BindCraft freedom to design additional interaction points of its own. Second, hotspot residues were required to be within 10 Å of one another (C_α_–C_α_ distance) to increase the chance that each binder will cover the entire hotspot. Third, each hotspot required a minimum of two hydrophobic amino acids, since burying hydrophobic side chains is a major contribution to binding free energy. Fourth, hotspot amino acids were required to be at least partly exposed on the protein surface (determined with the “surface” feature on ChimeraX) for accessibility. Fifth, hotspot amino acids were preferentially located on alpha-helices or beta-sheets to reduce entropic costs associated with immobilizing flexible loops upon binding. In total, 7 hotspots were selected out of 13 candidates (with 5 yielding promising binders), involving the coiled-coil domain, linker domain, SH2 domain and DNA-binding domain of STAT3 to create a collection of promising binders for areas that have been studied in relation to cancer progression, as well as surface regions that haven’t been studied as much to provide more future possibilities.

### 2.3 De novo design via the BindCraft pipeline

BindCraft’s default settings were modified to increase the penalties for designed binders that bind distally from input hotspot locations. The weights_con_inter parameter was raised from 1.0 to 2.0, to increase the importance of structural contacts between the binder and STAT3. We decreased inter_contact_distance, the maximum distance between the binder and STAT3, from 20 Å to 10 Å. The inter_contact_number parameter, which dictates the density of points of interaction between the binder and STAT3, was revised from 2 to 3. These alterations, along with other BindCraft parameter settings, are listed in Table 1. Cysteine was omitted from binder sequences to avoid folding complexities associated with requirements for correct disulfide bonds for binders to fold. An initial RFdiffusion/ProteinMPNN/AF2 [7] pipeline was used for hotspots 1–3, but it yielded no binders with ipTM above 0.30. These hotspots are considered preliminary trials and are not reported in this paper. All subsequent hotspots were designed with BindCraft, which automated the full binder design pipeline and is the source of all binders reported here.

**Table 1A.**
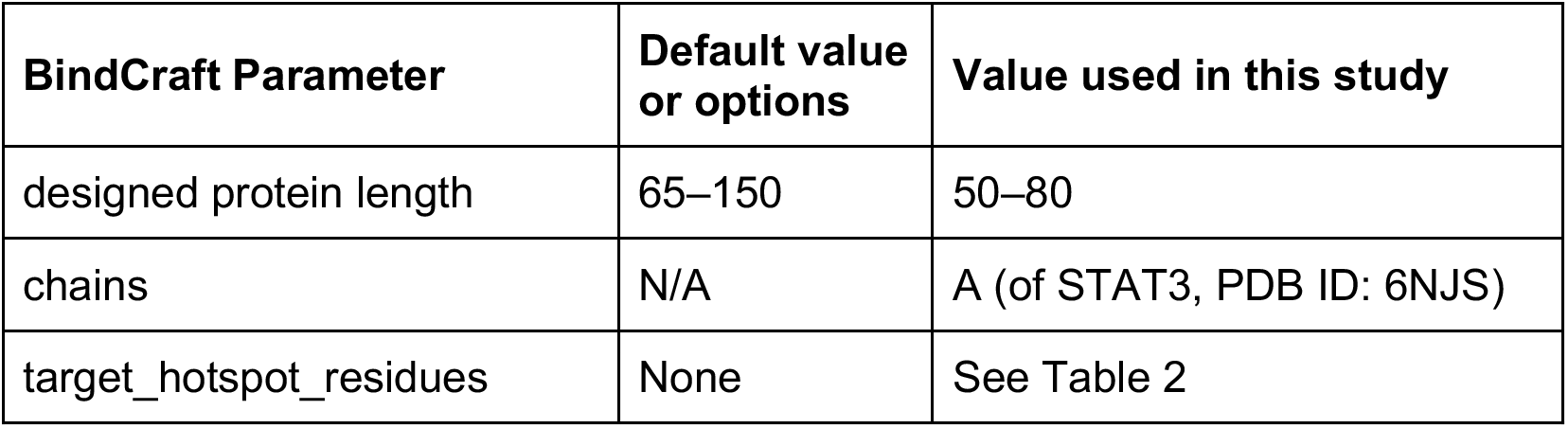
BindCraft Settings.

**Table 1B.**
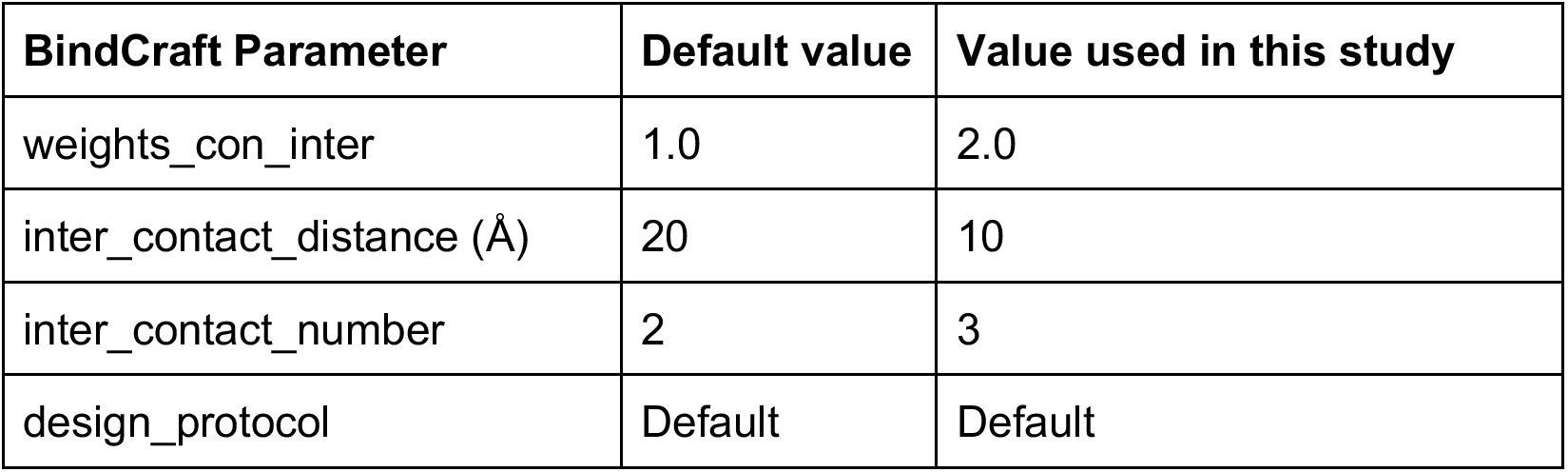
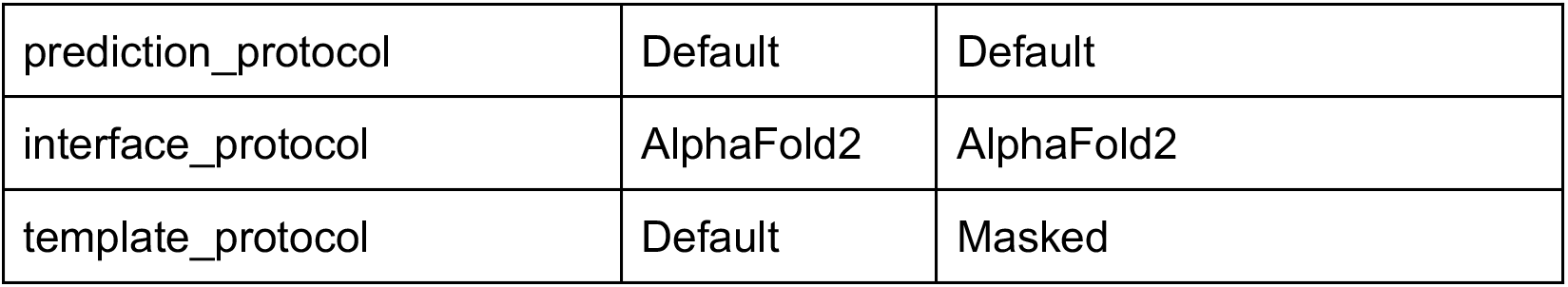
Advanced BindCraft Settings.

### 2.4 Additional scoring of designed binders with AlphaFold3

BindCraft incorporates AlphaFold2 (AF2) within its design pipeline. Replacing this module with AlphaFold3 (AF3) [8] would be technically challenging, particularly within Google Colab. However, AF3 is more advanced and regarded as more accurate than AF2 [9], so we used it as an downstream evaluator by manually submitting sequences to AFS.

We evaluated designed minibinders with BindCraft (AF2) ipTM values of 0.60 or greater using AlphaFold Server (AFS), which runs AF3. For each prediction, we recorded the interface predicted template modeling score (ipTM), predicted local distance difference test (pLDDT), and predicted aligned error (PAE). Designs were considered high confidence if they had an AF3 ipTM ≥ 0.75 and a pLDDT ≥ 90 for a grand majority of residues in the binder-target complex (the latter determined by visual inspection of theAF3-generated ribbon structure, which colors backbone segments by pLDDT range).

### 2.5 Residue numbering

Residue numbers quoted in this study follow “AlphaFold Server (AFS) numbering.” AFS output sequences always begin at 1. The length of STAT3 in the 6NJS crystal structure file (available from the protein data bank (PDB) at https://www.rcsb.org/structure/6NJS) is 562 amino acids. The AFS numbering scheme for STAT3 from 6NJS is therefore1 to 562. In the 6NJS PDB file, the first amino acid in the sequence of STAT3 is numbered 127. In the 6NJS structure downloaded from the PDB itself, the first amino acid is assigned #127. Therefore, there is a constant offset between the two numbering systems of 126: aa_number_PDB_6NJS_ = aa_number_AFS_6NJS_ +126. A STAT3 sequence numbering guide that compares the two numbering schemes is provided as Figure 6.

### 2.6 Structural visualization

High-confidence binders were visualized with ChimeraX. The target protein, hotspot and binder were all colored differently and hotspot residues were labeled (Figures 3–4). Both ribbon and sphere modes were applied to visualize structures and binder interactions with STAT3. Figure 4 depicts two binders simultaneously docked with STAT3. To create Figure 4, we did not input the sequences of STAT3 and both minibinders into AF3 for a ternary complex evaluation, as the tool will distort folds and binding locations to “create” a ternary complex. Instead, we opened the two separate AF3 models of each individual minibinder docked with STAT3 with ChimeraX, then performed a structural alignment.

**Figure 3.**
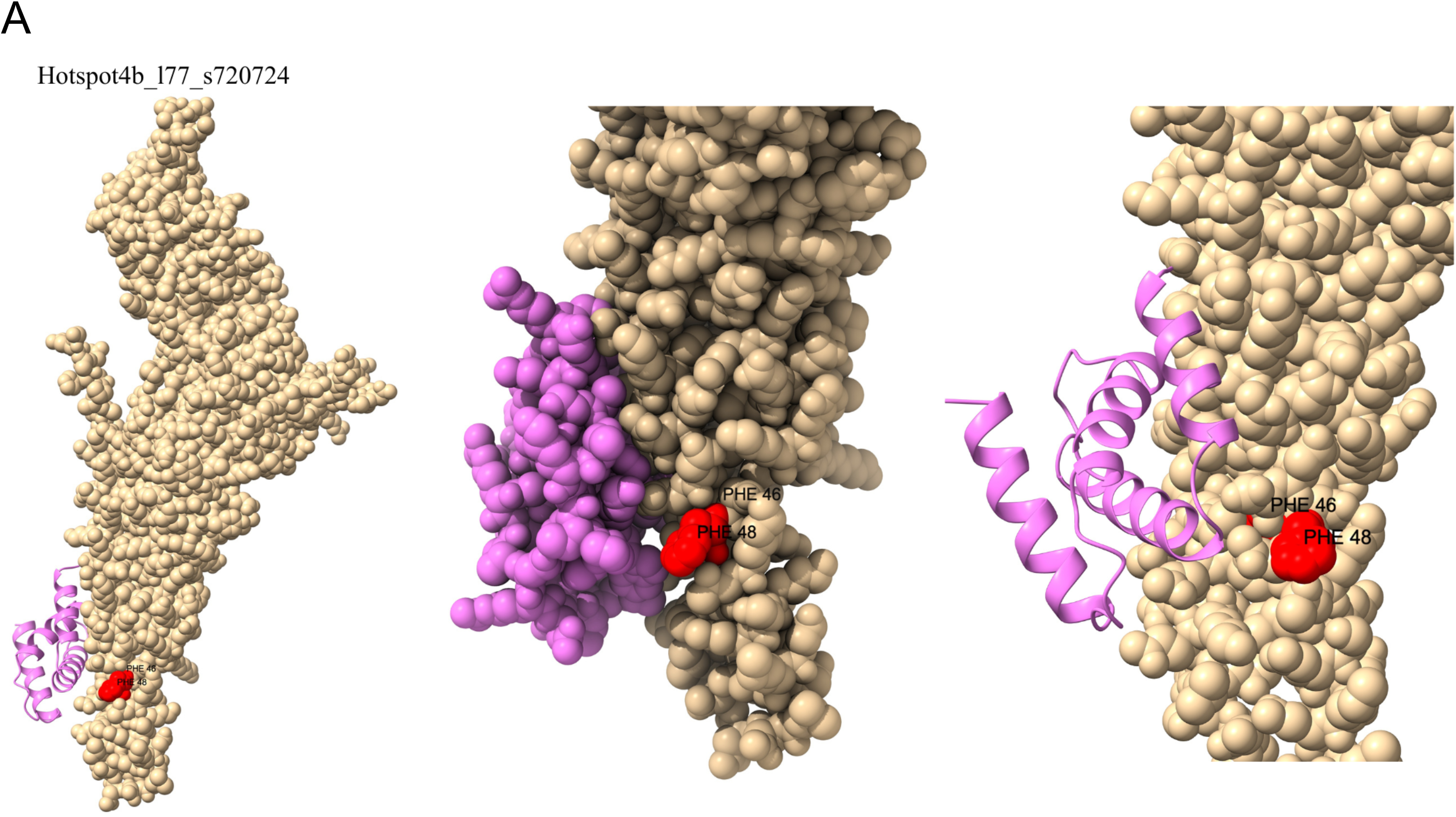
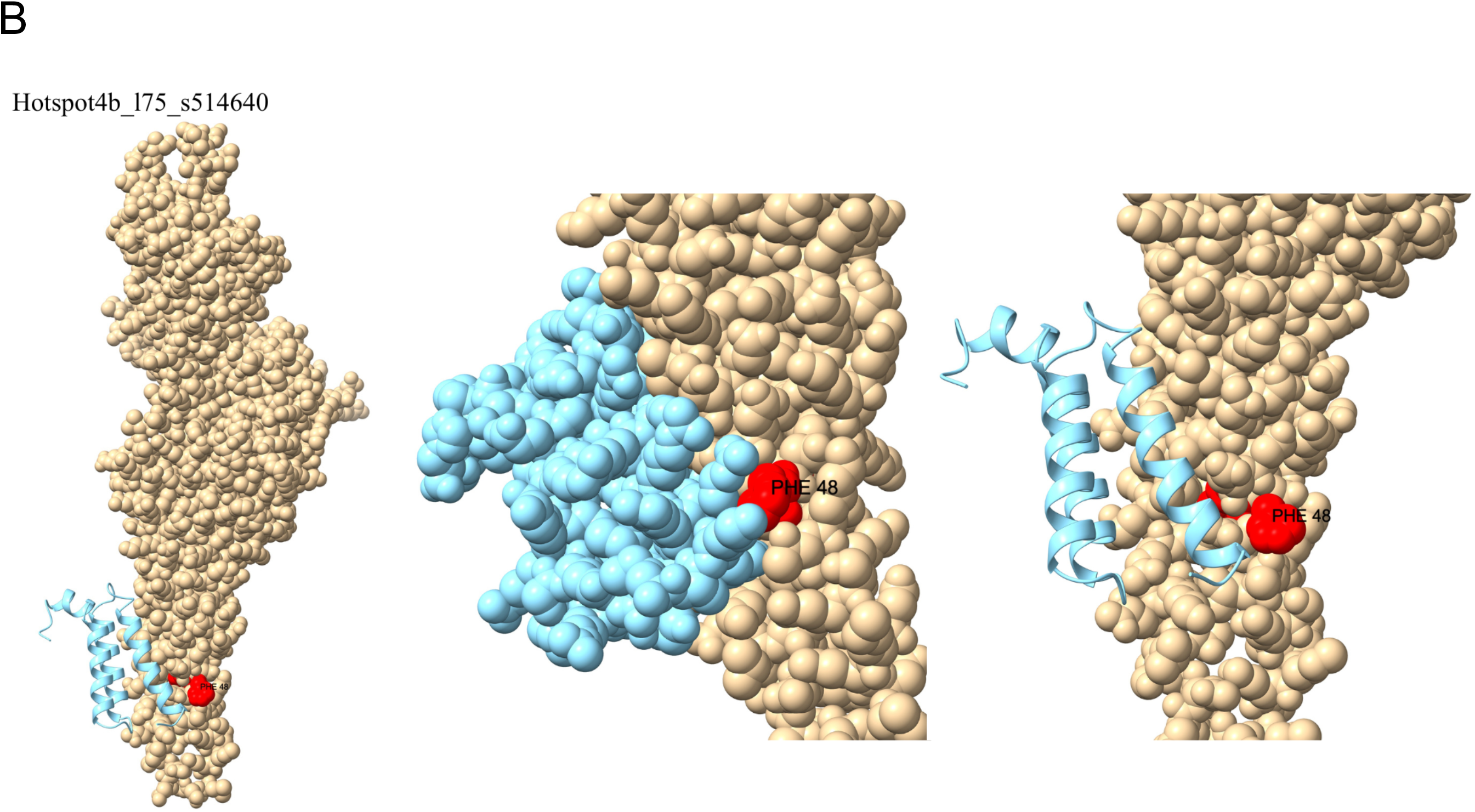
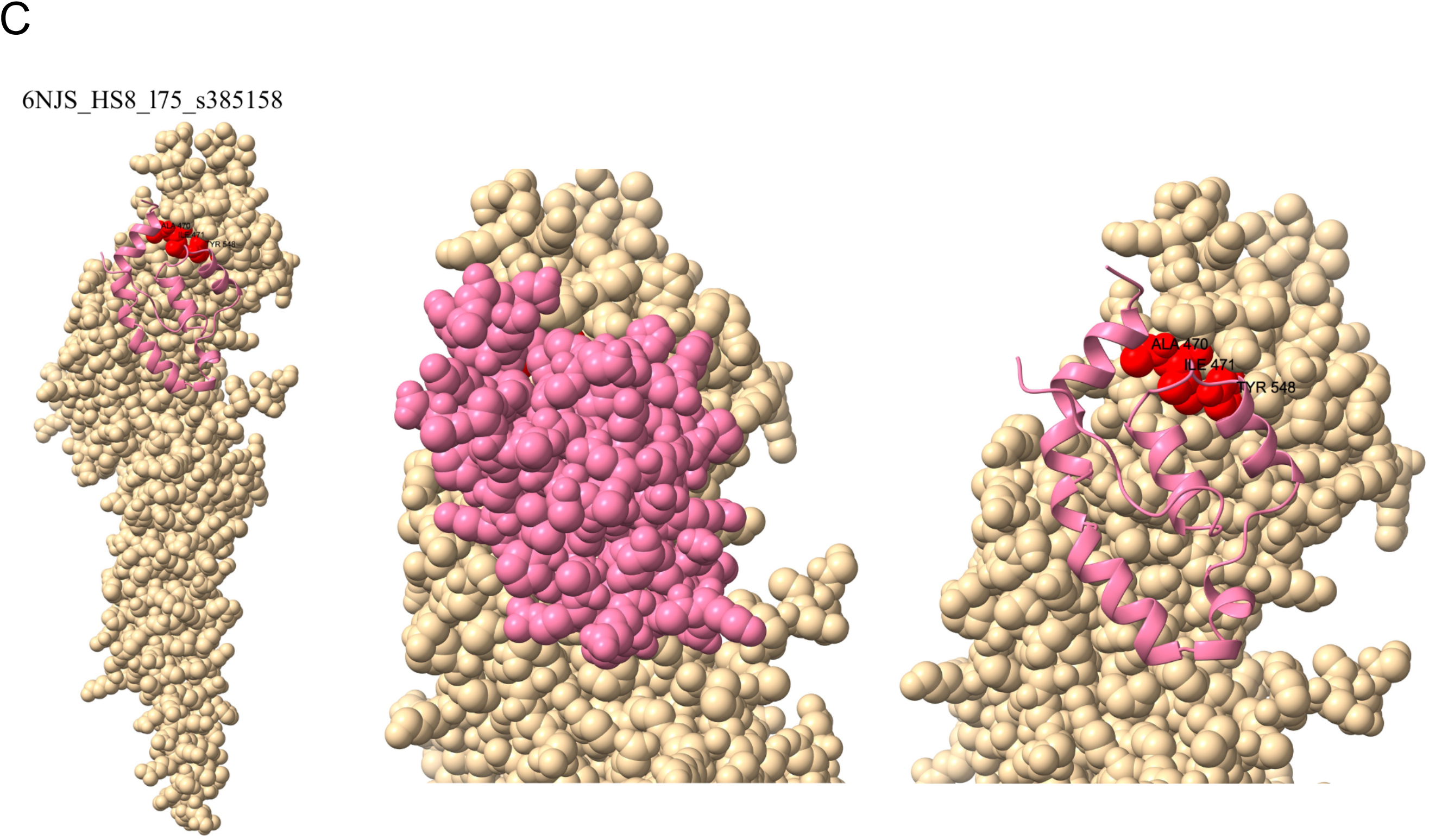
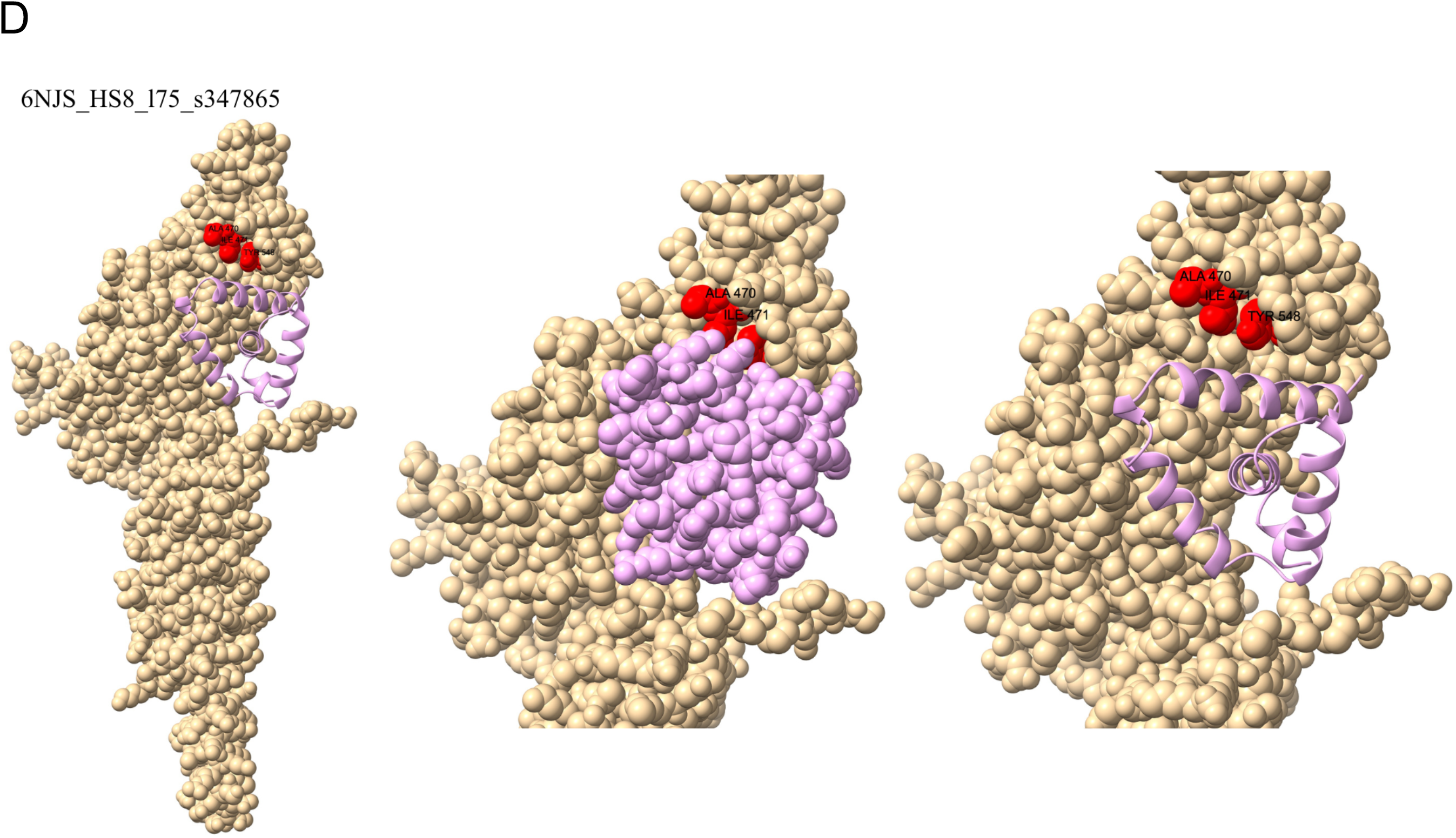
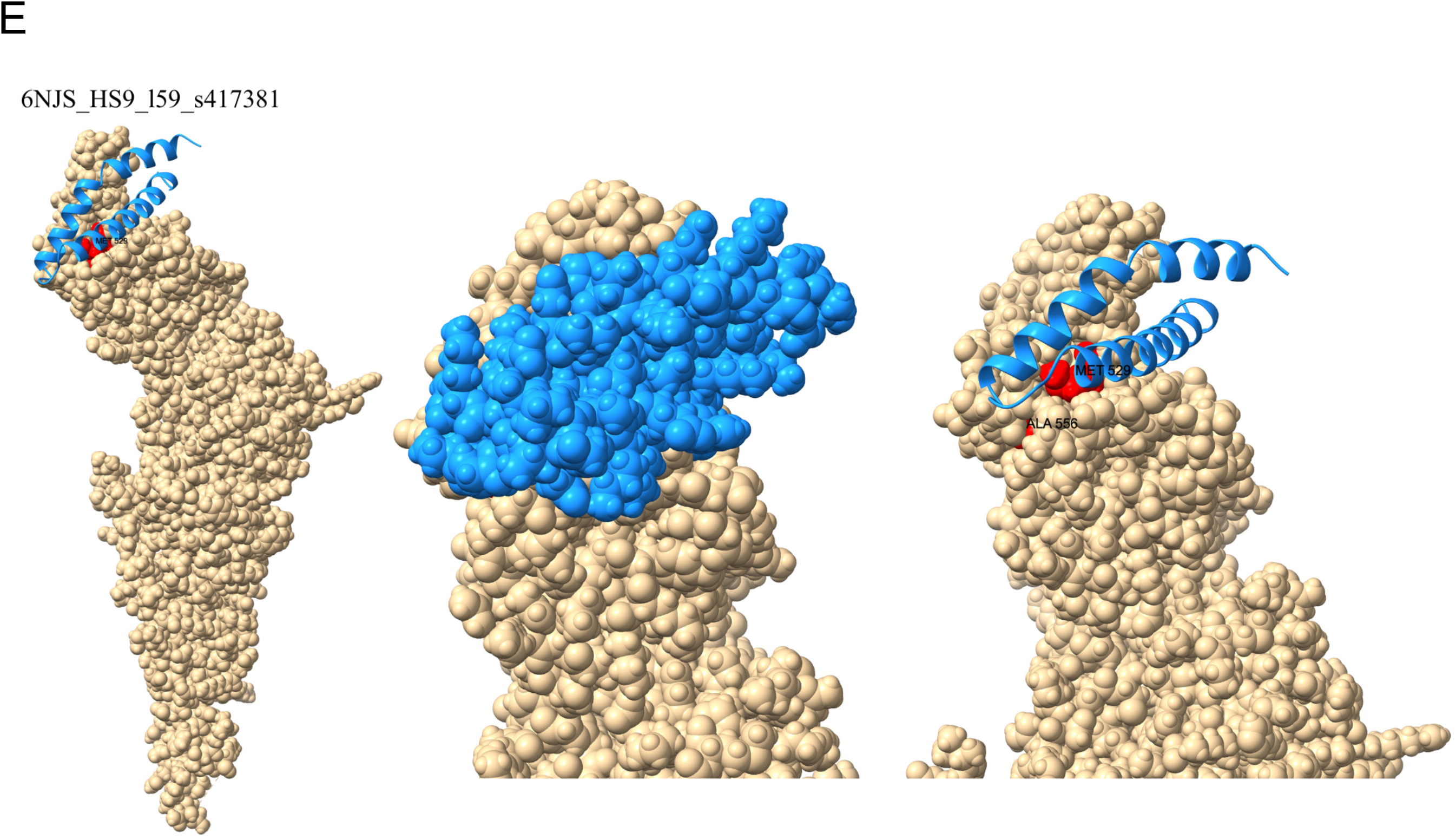
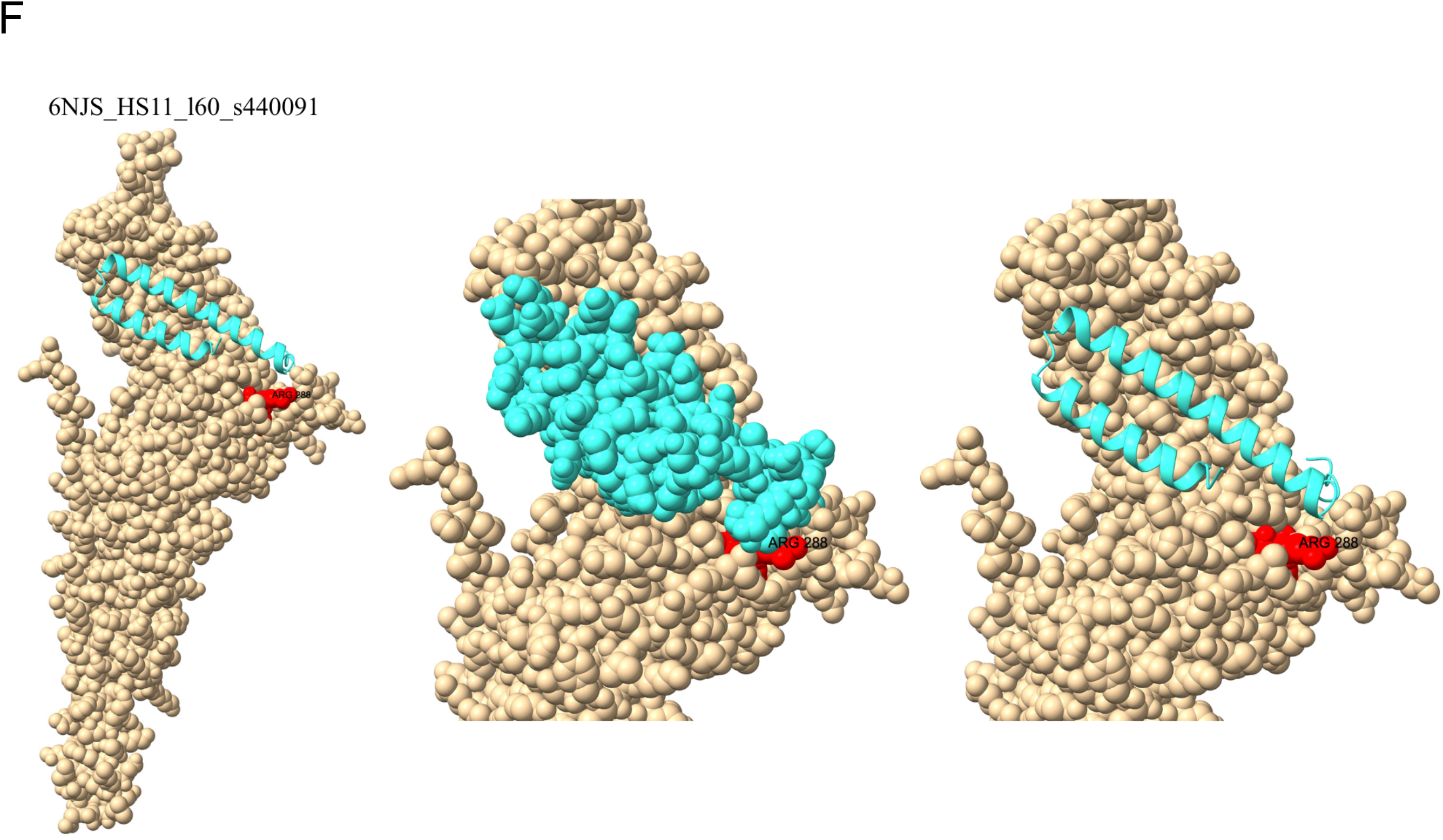
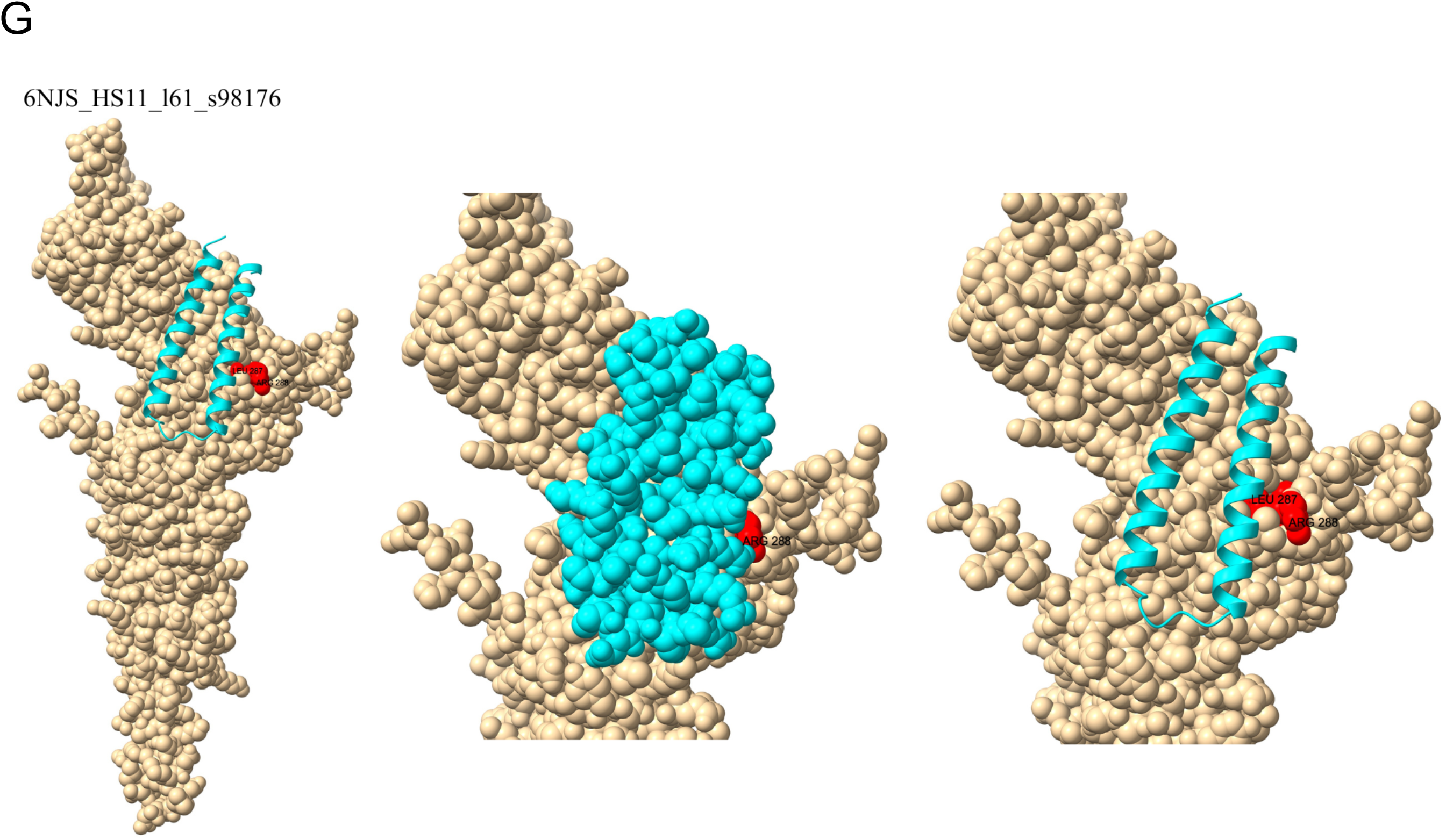
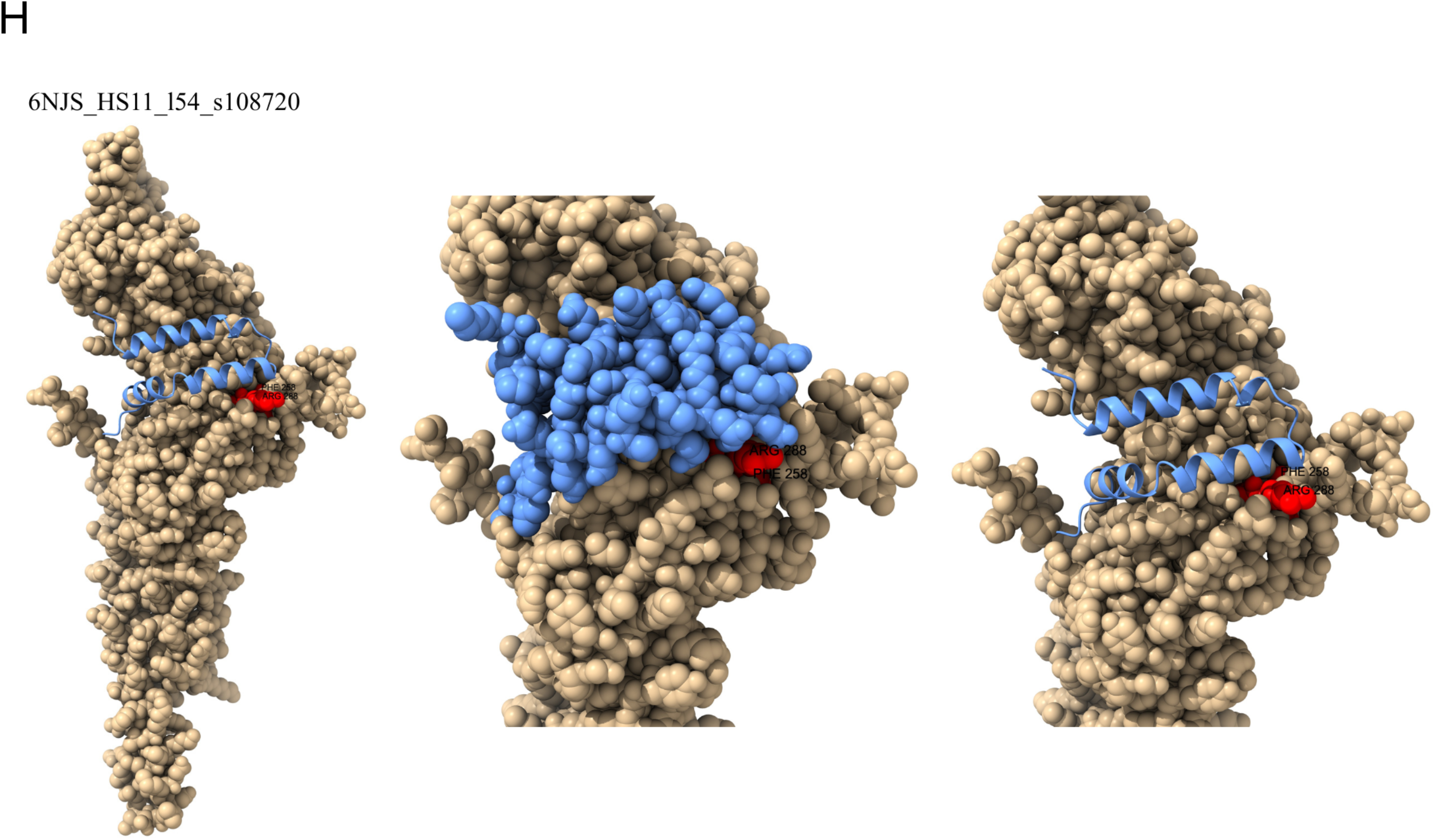
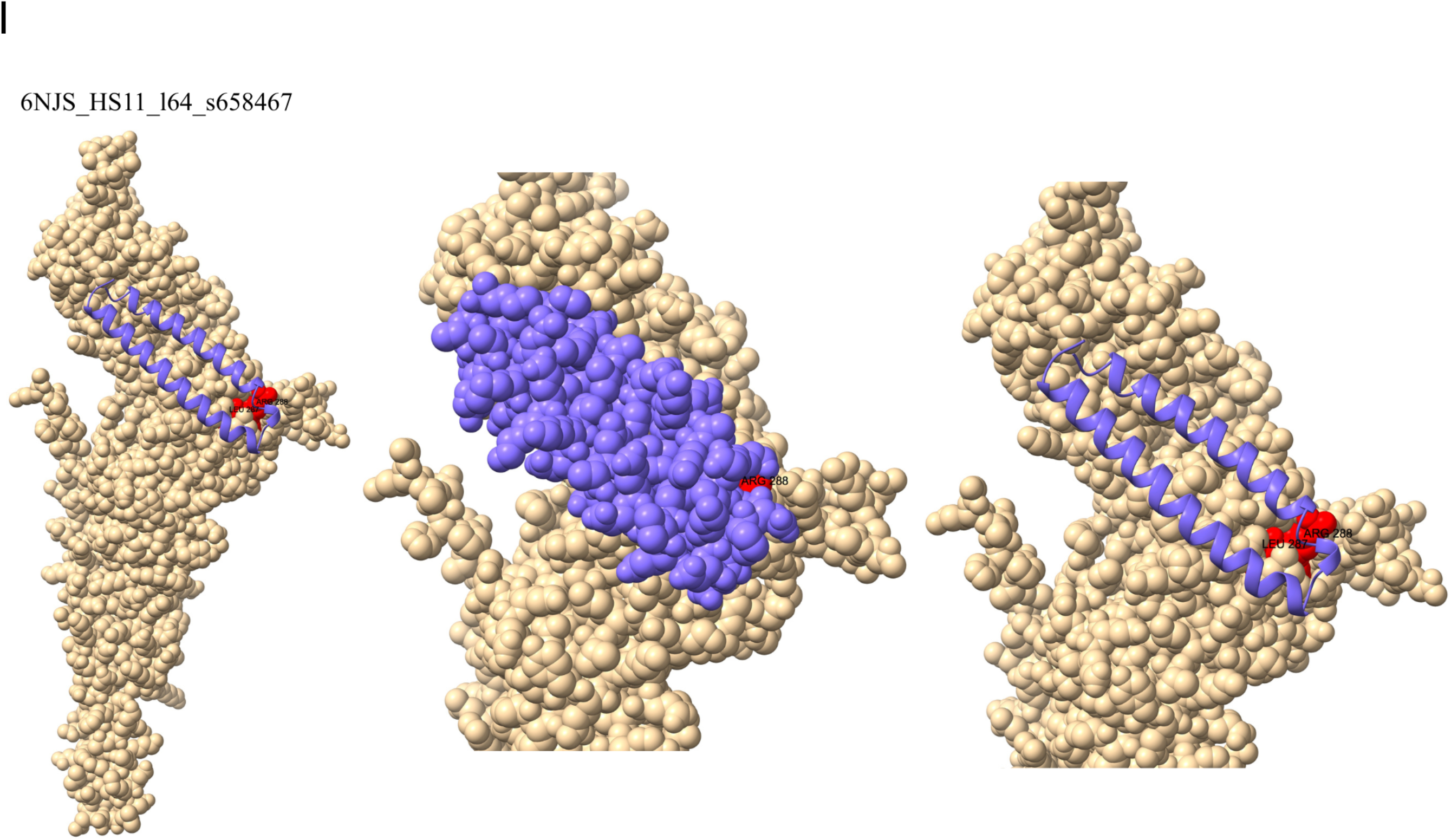
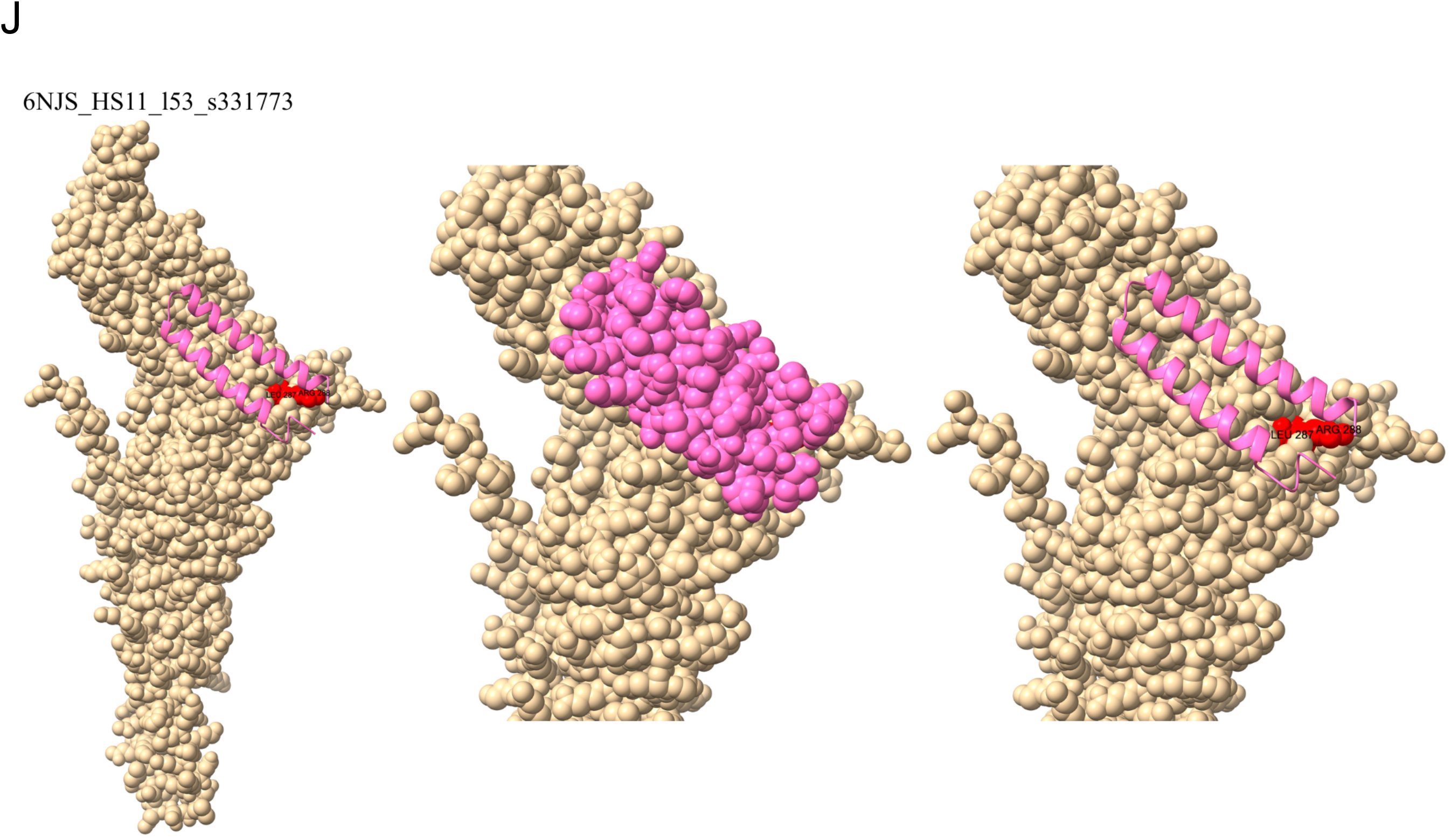
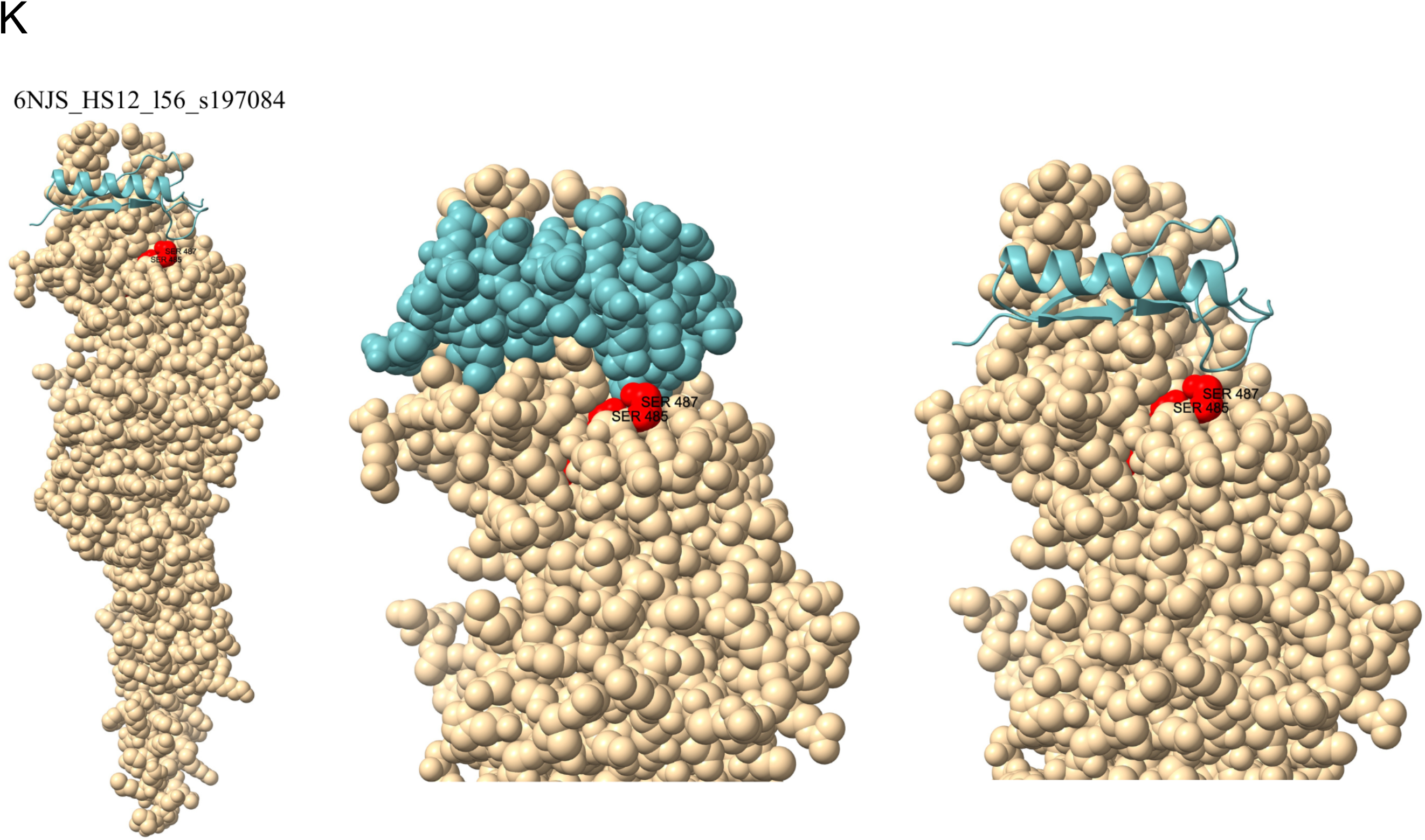
(Panels A–K spanning 11 pages) ChimeraX renderings of AlphaFold3 predictions of each high-confidence designed minibinder’s interaction with STAT3 (beige space-filling model). For each panel: Left, predicted docking location of the minibinder (colored ribbon) on STAT3 (whole protein shown). Center, detailed view of the predicted binding interaction with both chains presented as space-filling models. Right, detailed view of the predicted binding location with the minibinder shown in ribbon form. Red spheres mark the input hotspot amino acids for each binder’s design run; those not completely covered by a minibinder in each image are labeled (AFS numbering scheme). Binder names are provided at top-left of every panel. In panel H (6NJS_HS11_l54_s108720) Phe 258 is largely hidden by Arg 288.

**Figure 4.**
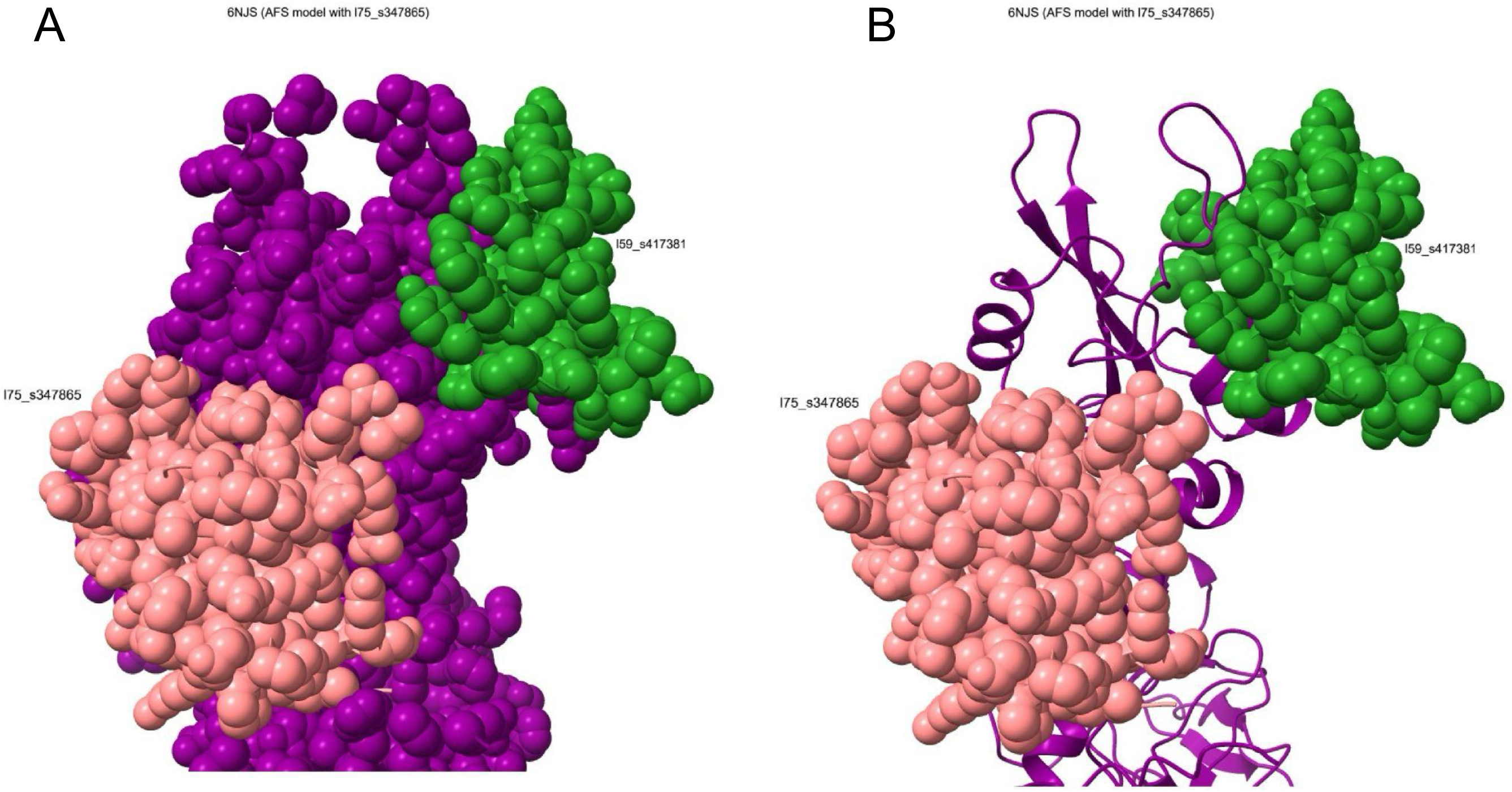
Composite view of minibinders l75_s347865 (pink), l59_417381 (green) and STAT3 (purple) bound simultaneously to non-overlapping faces of STAT3, illustrating their potential as a sandwich-assay pair. A, detailed view of the two binder-STAT3 ternary complex; all chains shown as atomic spheres. B, with STAT3 shown in ribbon form. See section 2.6 for a description of how this figure was generated.

### 2.7 Sequence novelty

Every high-confidence binder was subjected to a BLASTp search to assess its sequence novelty. The BLASTp settings employed were: non-redundant (NR) protein database, E-value cutoff threshold of 10, seed word size of 5, and BLOSUM62 matrix.

### 2.8 Assessment of binder-hotspot overlap

We adopted a common definition for a protein-protein interface of amino acids within 4.5 Å of any non-hydrogen atom of a designed binder (Table 3) [10, 11]. To identify interface amino acids of STAT3, the command “contacts (/A & ∼@H) restrict (/B & ∼@H) distance 4.5 log true” was run in ChimeraX. We compared the input hotspot residues entered in the design pipeline for each binder against its list of interface residues. We also compared the targeted domain of STAT3 with the domain of STAT3 that interacts with each binder, according to AF3. Additionally, we visually assessed the degree of “overlap” or coverage of each input hotspot by its corresponding designed minibinder using “spheres” mode in ChimeraX. For the purposes of this study, overlap is defined as STAT3 residue whose surface appears to be ≥ 50% covered by a binder, according to the AF3 output structure. Results are listed in Table 5.

## 3. Results

### 3.1 Results overview and pipeline yield

Across 7 hotspots, 17 BindCraft campaigns yielded 139 recovered sequence designs, 98 of which had a pipeline (AF2) ipTM ≥ 0.60. These 98 were subjected to evaluation of folding and binding to STAT3 by AlphaFold Server (Table 3). Eleven designs from 5 of the 7 design hotspots met the threshold of ipTM ≥ 0.75 (Tables 2 and 3). Hotspot 11 was the most productive (5 passing designs from 3 campaigns), followed by hotspots 4B and 8 (2 each) and hotspots 9 and 12 (1 each). Hotspots 4A and 7 produced no designs that met the AF3 thresholds. BindCraft ipTM values range from 0.81–0.94, AF3 ipTM values range from 0.75–0.93. Binders are shown in Table 3 with their respective ipTM and sequences. An overview of pipeline yield is shown in Table 2.

**Table 2.**
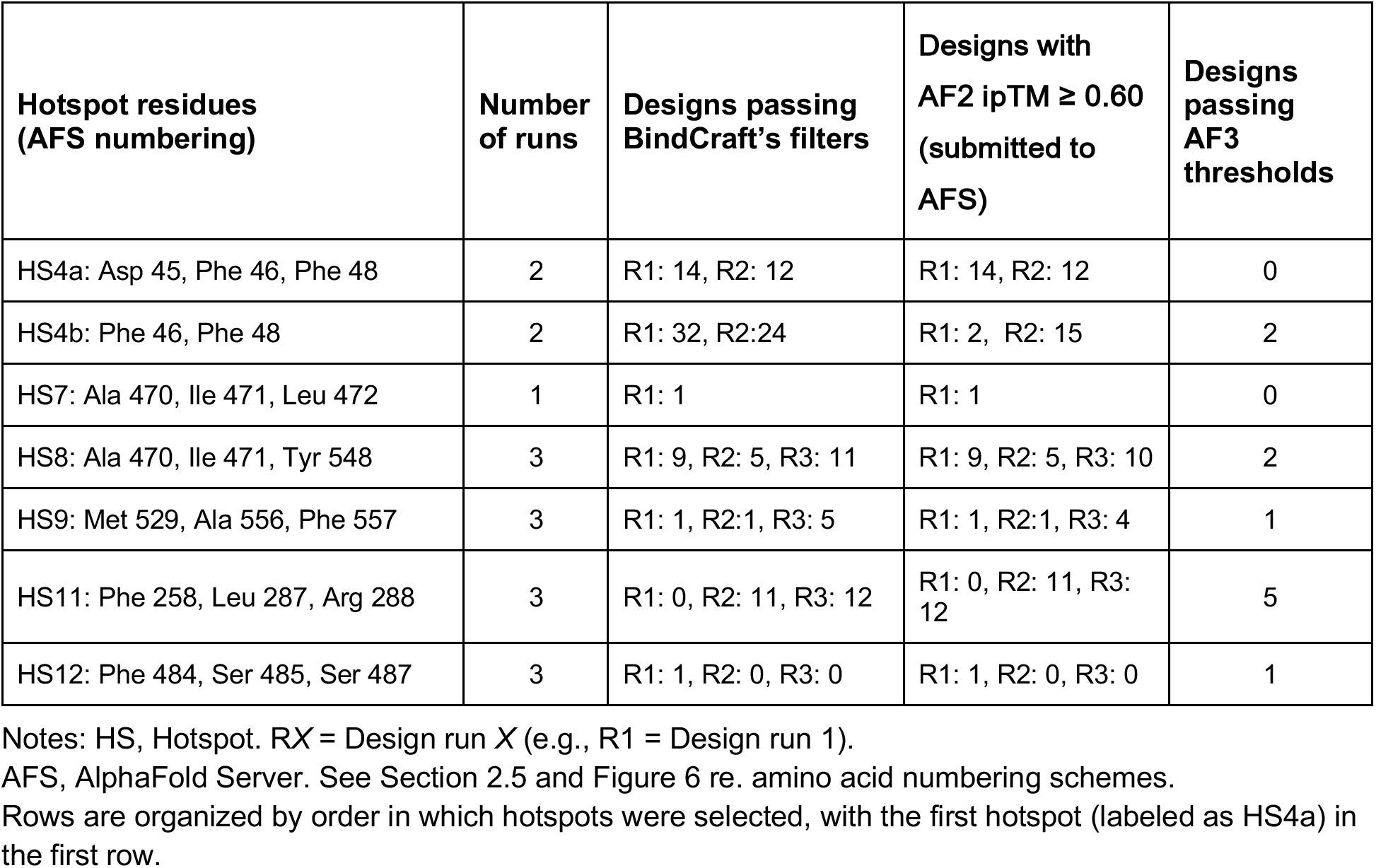
BindCraft input hotspots and design yield statistics.

**Table 3.**
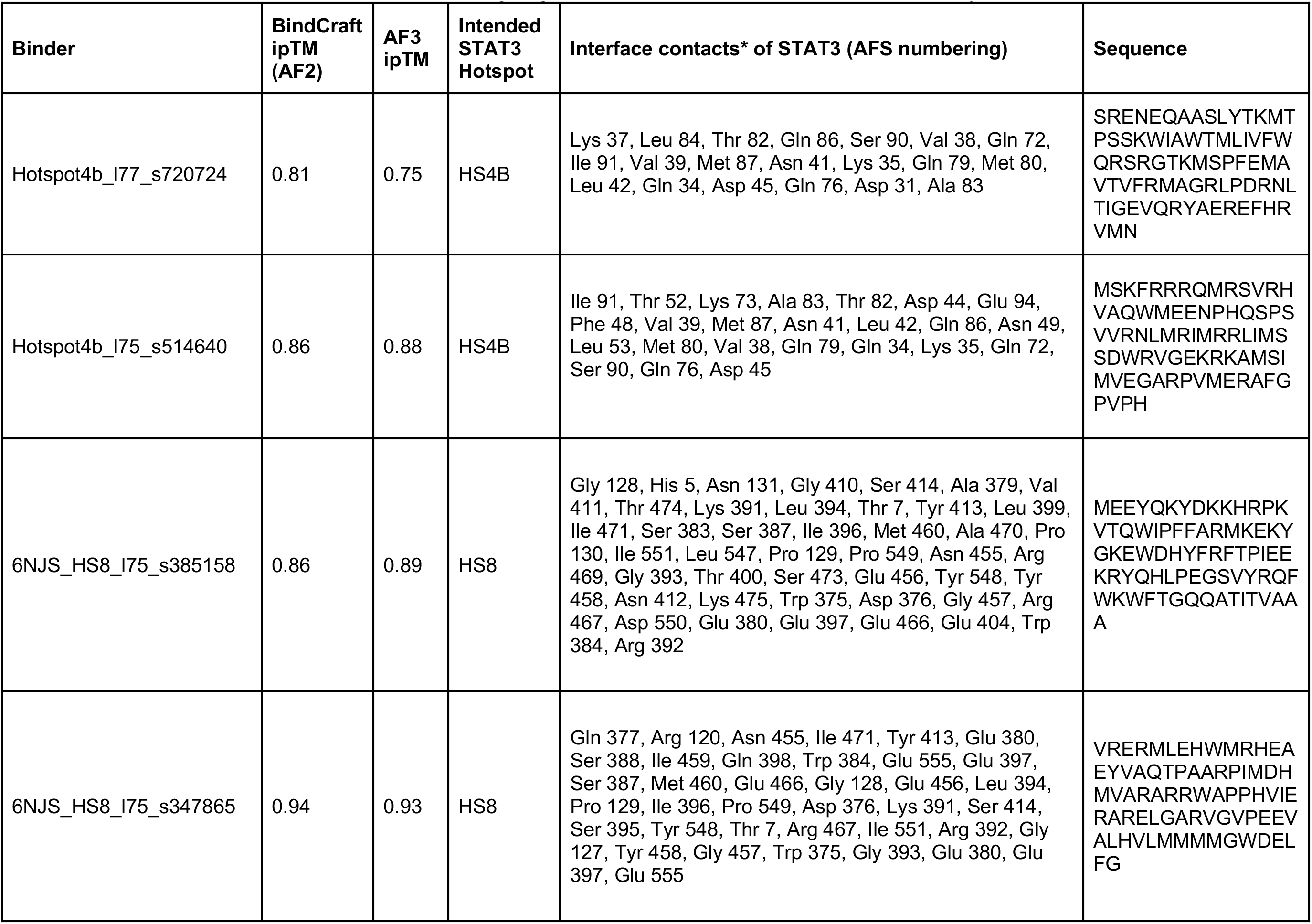
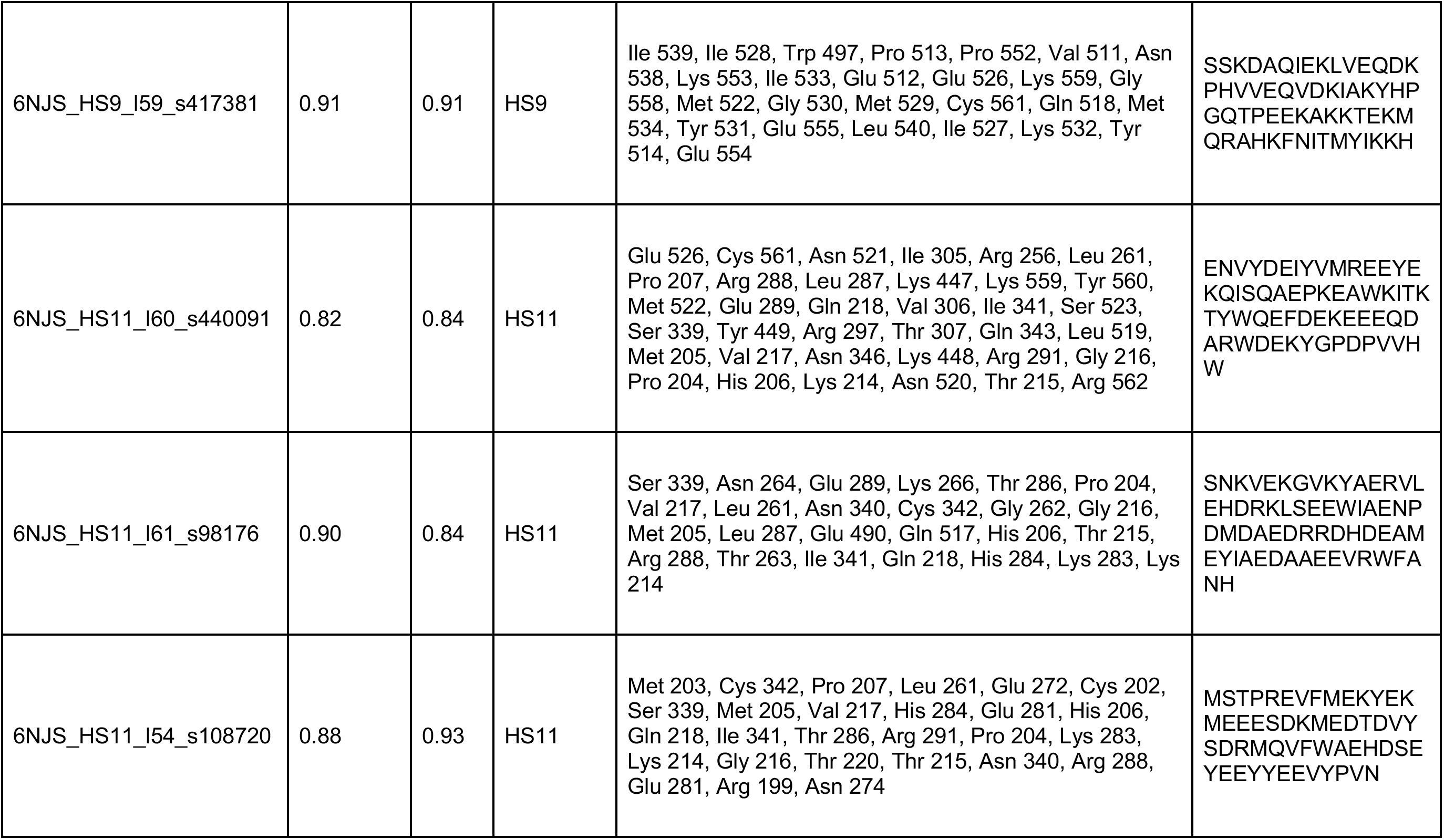
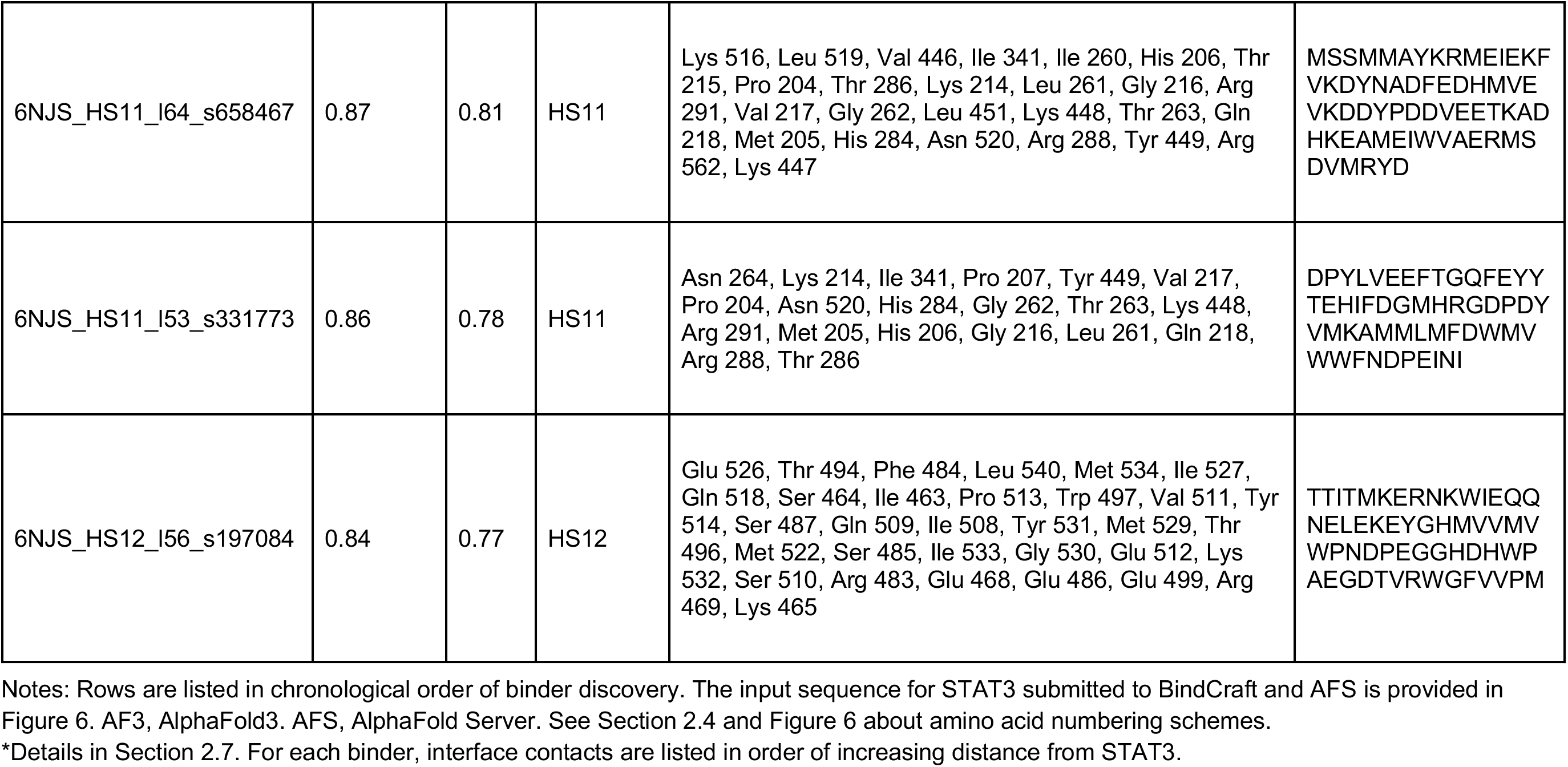
De novo candidate STAT3 minibinder designs generated and evaluated in silico in this study.

### 3.2 Structural modeling of top candidate binders

Figure 3 depicts AlphaFold3 structural models of all eleven top binders interacting with STAT3. The binder, STAT3 and the intended hotspot are all distinctly colored for identification. The STAT3 hotspot residues input into BindCraft are all colored red (and labeled if visible) to highlight the degree of overlap between each top binder and its input STAT3 hotspot residues. Van der Waals atomic sphere view helps visualize the interactions between STAT3 and each binder, while ribbon view clarifies binder folds.

Binders 6NJS_HS8_l75_s347865 and 6NJS_HS9_l59_s417381 are additionally depicted bound to STAT3 simultaneously, as they do not overlap each other and are therefore candidates for a two-site sandwich assay (Figure 4). These two minibinders are not the only sandwich assay candidates among the set; they were merely selected as an illustrative example.

In the figures, labeled residue numbers correspond to the AlphaFold Server numbering scheme.

### 3.3 Confidence assessment via predicted aligned error

To further evaluate the interfaces the 11 top-ranked minibinder candidates form with STAT3, the Predicted Aligned Error (PAE) plots generated by AFS for each independent run were inspected (Figure 5). The off-diagonal rectangular blocks representing the binder-target interface show low predicted error for all binders (more green, less white area). This signals that AlphaFold is confident in the mutual match of each protein with the other at the interface.

**Figure 5.**
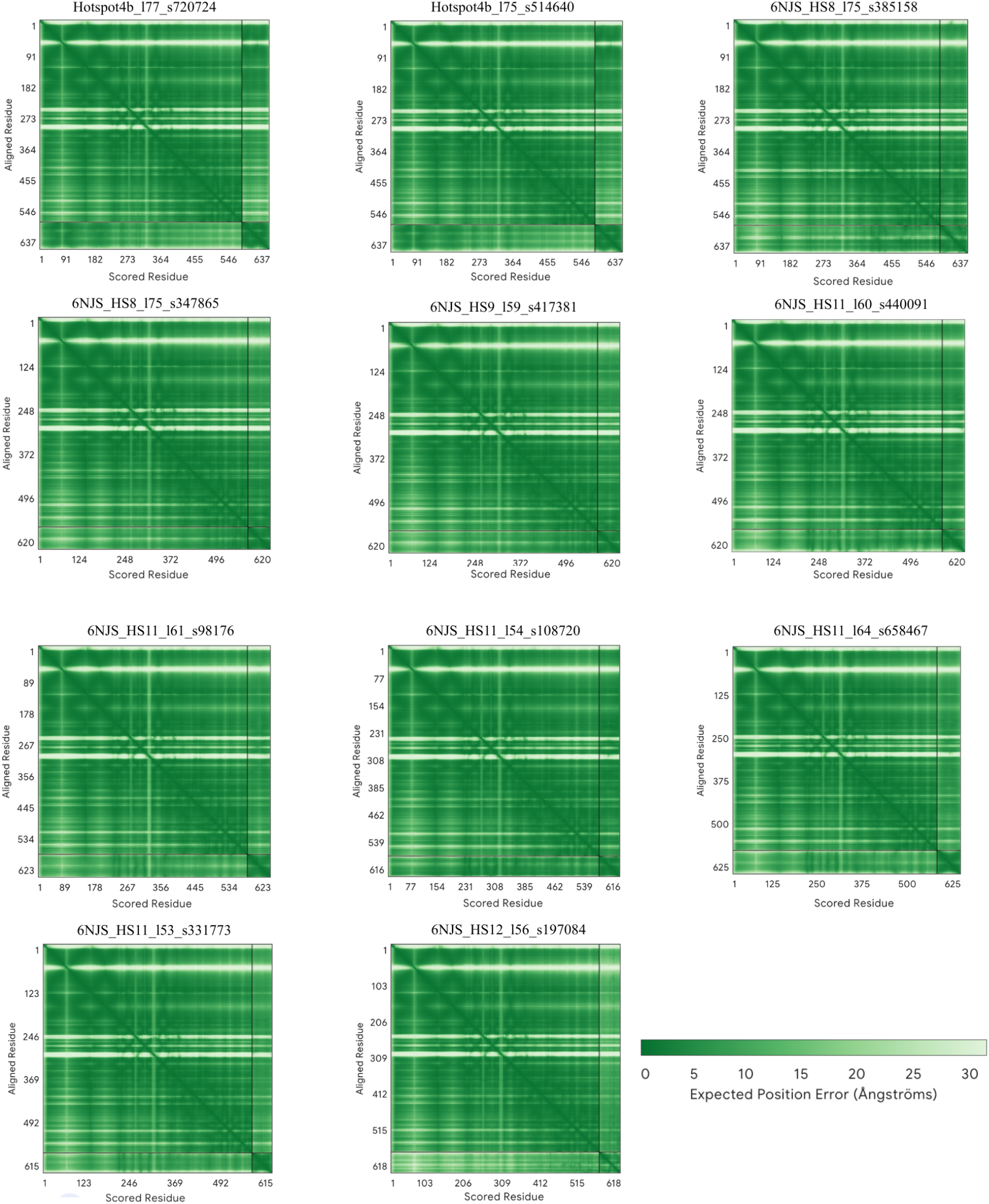
Predicted Aligned Error (PAE) plots from AlphaFold Server (AF3) evaluation of 11 high-confidence STAT3 binders. Each heatmap plots the expected position error (Å) for every pair of residues in the modeled complex. Deeper green coloring (lower error, see legend) in the top-right and bottom-left off-diagonal rectangular blocks, which correspond to the binder-target interface indicates confident, well-defined relative positioning between binder and target chains. Panels are labeled with binder identity.

**Figure 6.**
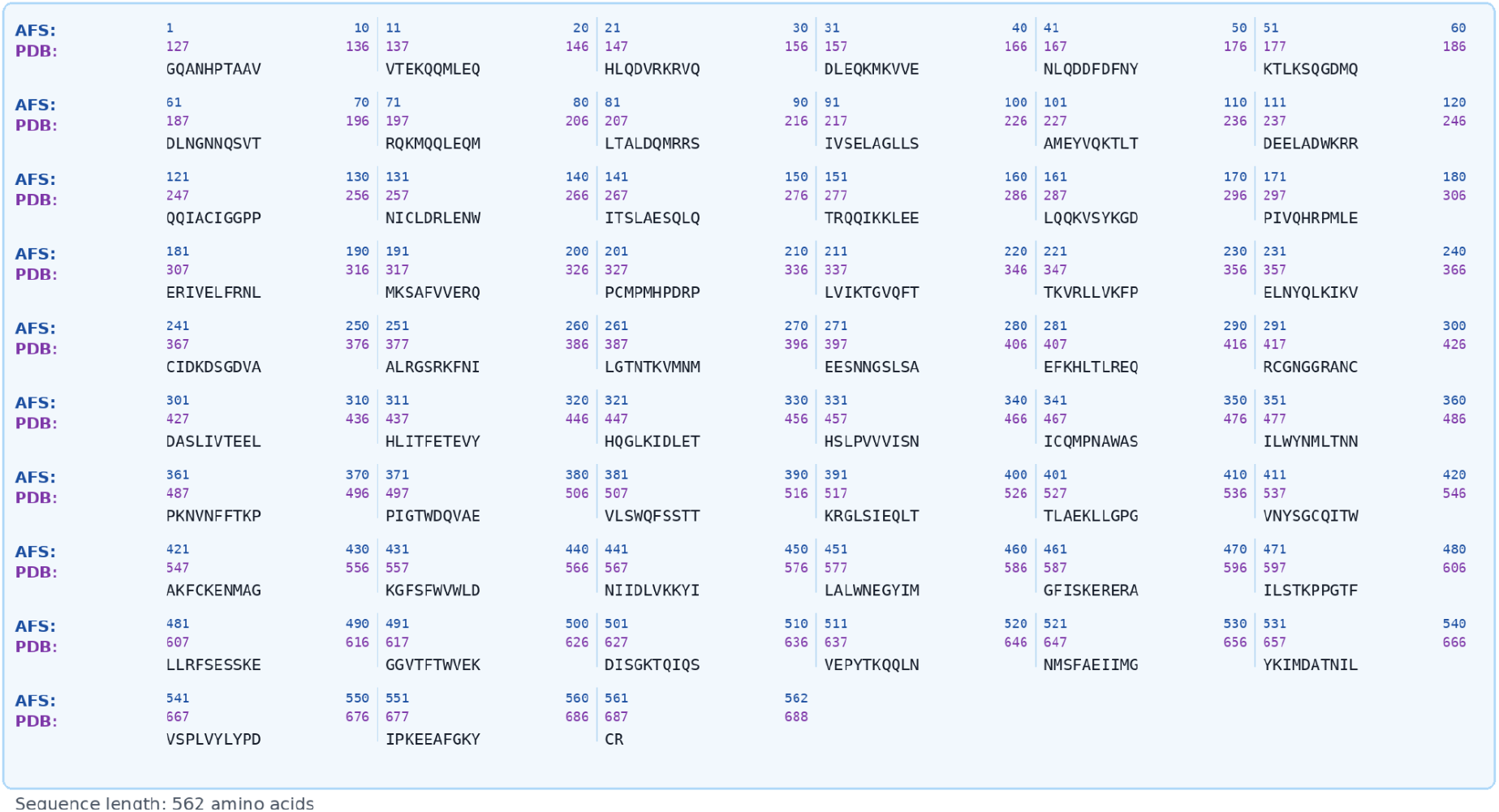
The STAT3 sequence from the 6NJS PDB entry input to BindCraft and AFS, shown with the AFS and 6NJS PDB numbering schemes. Amino acid number in the 6NJS PDB numbering scheme = Amino acid number in the AFS scheme + 126. Sequence text for copying: >STAT3 (6NJS) GQANHPTAAVVTEKQQMLEQHLQDVRKRVQDLEQKMKVVENLQDDFDFNYKTLKSQGDMQDLNGNNQ SVTRQKMQQLEQMLTALDQMRRSIVSELAGLLSAMEYVQKTLTDEELADWKRRQQIACIGGPPNICLDRL ENWITSLAESQLQTRQQIKKLEELQQKVSYKGDPIVQHRPMLEERIVELFRNLMKSAFVVERQPCMPMHP DRPLVIKTGVQFTTKVRLLVKFPELNYQLKIKVCIDKDSGDVAALRGSRKFNILGTNTKVMNMEESNNGSL SAEFKHLTLREQRCGNGGRANCDASLIVTEELHLITFETEVYHQGLKIDLETHSLPVVVISNICQMPNAWA SILWYNMLTNNPKNVNFFTKPPIGTWDQVAEVLSWQFSSTTKRGLSIEQLTTLAEKLLGPGVNYSGCQIT WAKFCKENMAGKGFSFWVWLDNIIDLVKKYILALWNEGYIMGFISKERERAILSTKPPGTFLLRFSESSKE GGVTFTWVEKDISGKTQIQSVEPYTKQQLNNMSFAEIIMGYKIMDATNILVSPLVYLYPDIPKEEAFGKYCR

### 3.4 BLASTp search results

NCBI BLASTp searches [12] of all 11 designed binder sequences indicated no matches to any known proteins for 6 of the binders and only very low-significance, coincidental matches for the other 5 (using a threshold E-value for statistical significance of sequence matches of 10^-5^ [13]). The low-significance matches are classified as either bacterial or hypothetical proteins, and have no relation to STAT family or known mammalian sequences. These results suggest that all designed binders are novel sequences, not chimeras of already-known proteins. These results are shown in Table 4.

**Table 4.**
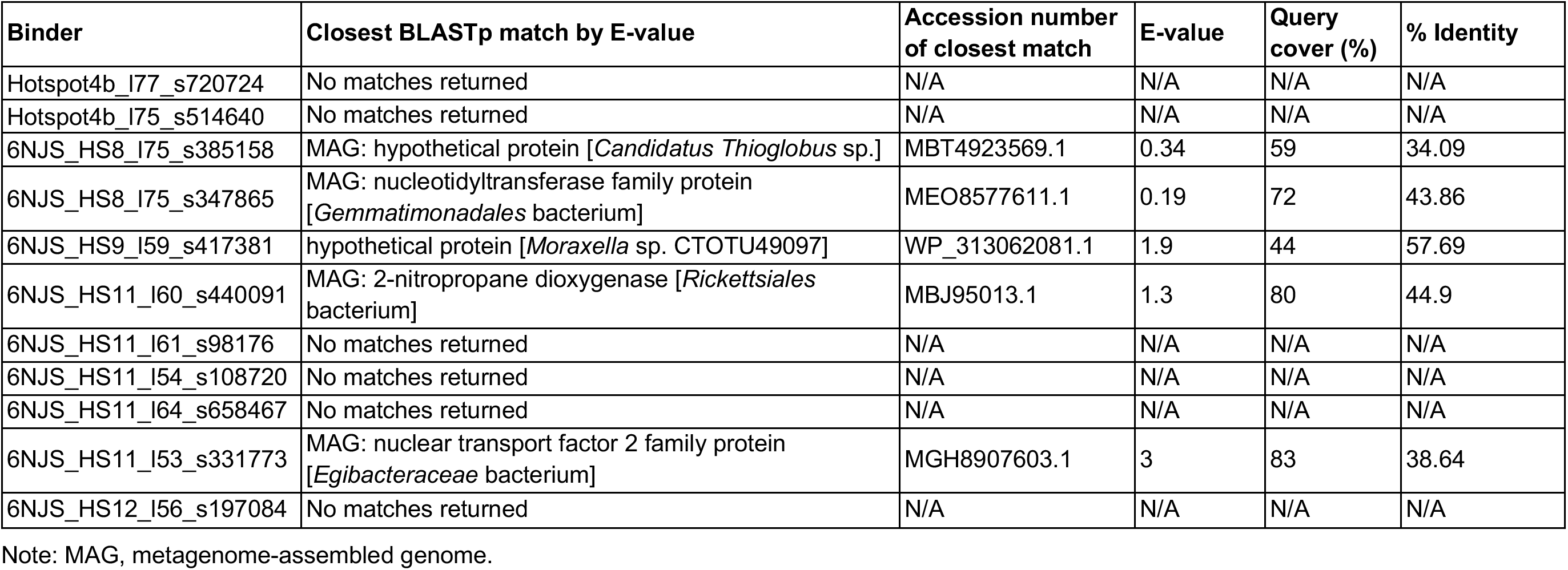
BLASTp search results for 11 high-confidence minibinders.

### 3.5 Overlap between binders and intended hotspot

We compared the AF3-predicted binding site of each of the 11 top binders with its input hotspot residues and targeted interaction domain (Table 5). Two measures, defined in the notes for Table 5, were used. Ten of the 11 binders had at least one hotspot residue within 4.5 Å of a STAT3 residue. For two minibinders (6NJS_HS8_l75_s385158 and 6NJS_HS12_l56_s197084), all hotspot residues are also interface residues, while eight minibinders have one or two hotspot residues among their interface residues. Only Hotspot4b_l77_s720724 has no hotspot residues listed among its interface residues. By visual inspection of overlap (coverage), nine of the 11 binders cover at least one hotspot residue. Four binders cover all their input hotspot residues (6NJS_HS8_l75_s385158, 6NJS_HS9_l59_s417381, 6NJS_HS11_l64_s658467 and 6NJS_HS11_l53_s331773), five cover some, and two cover none (Hotspot4b_l77_s720724 and 6NJS_HS12_l56_s197084).

**Table 5.**
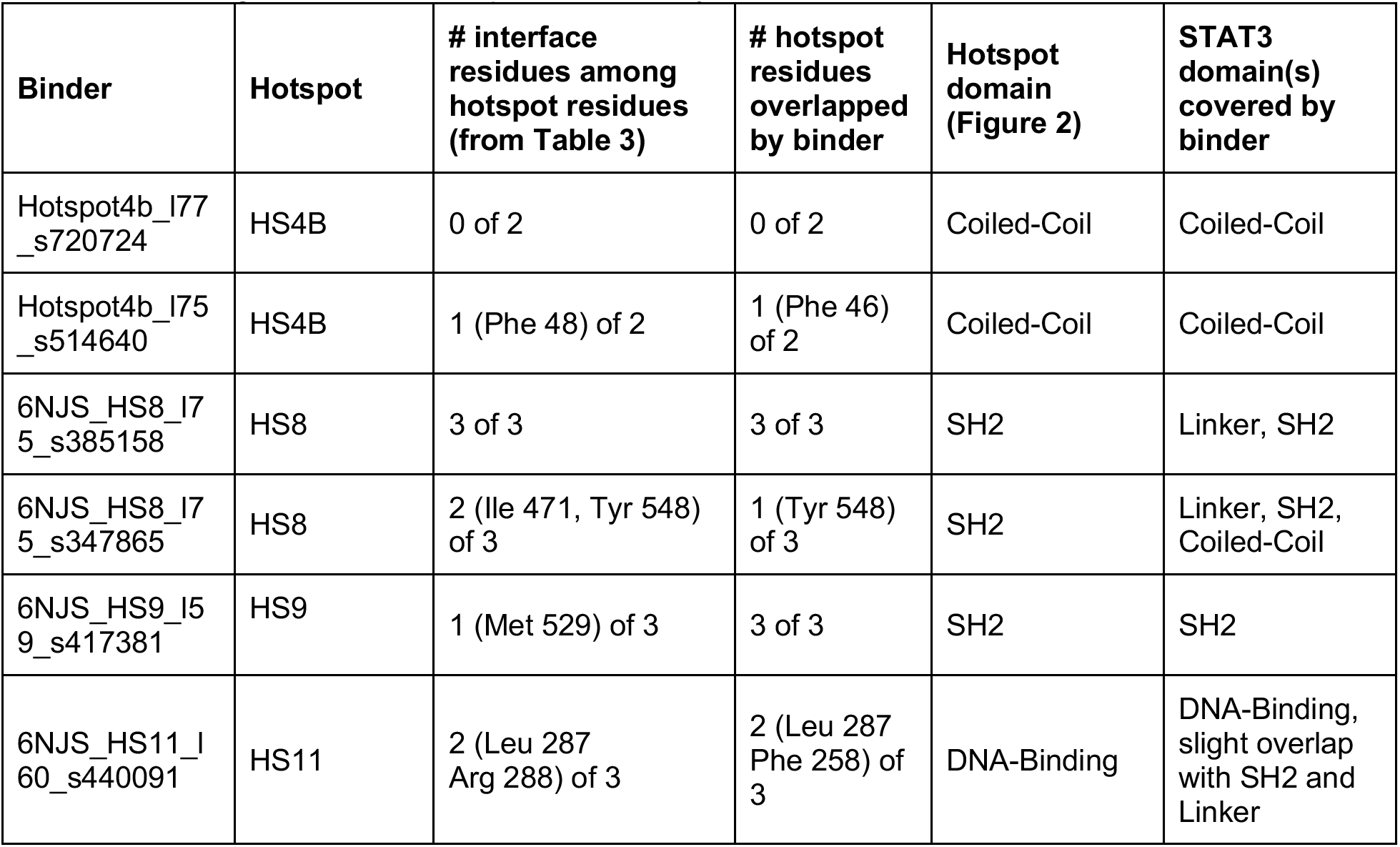
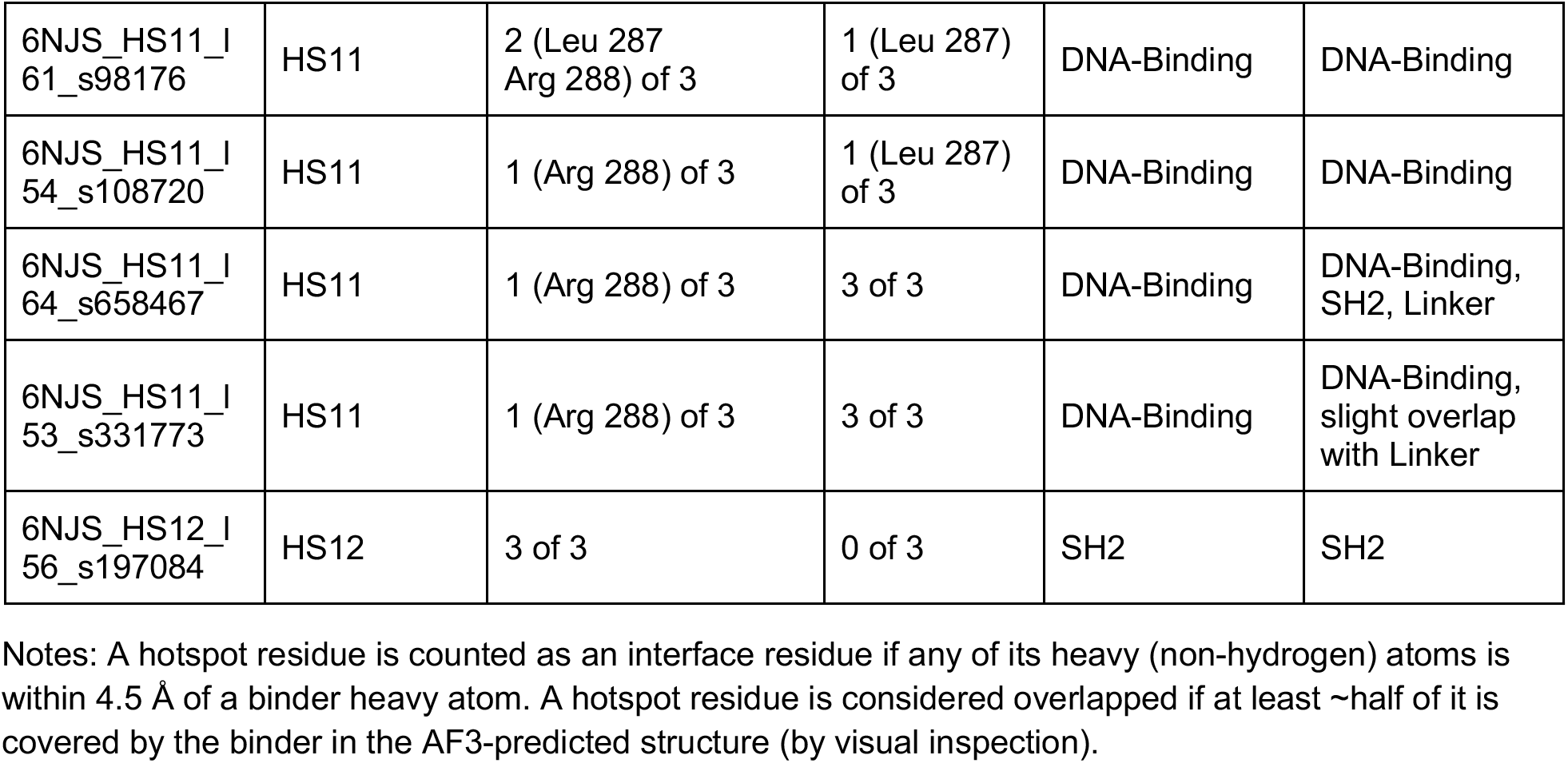
Coverage of STAT3 hotspot residues by minibinders.

All 11 binders cover their intended domain. Six dock only with the intended domain (both HS4B binders, the HS9 binder, the HS12 binder and two HS11 binders). The other five also cover adjacent domains: both HS8 binders also contact the linker domain (one also into the coiled-coil domain), and three HS11 binders cover part of the SH2 and/or linker domains.

## 4. Discussion

### 4.1 Biological significance of the targeted regions

We chose hotspots by structural criteria, but we made sure to consider biological significance, as the ultimate purpose is to provide binders with the potential to modulate or interfere with STAT3’s involvement in cancer pathogenesis or other diseases. Table 6 lists each STAT3 hotspot in both the AFS and the 6NJS PDB numbering schemes, as well as the domain’s known or assigned function. While not every minibinder covers its entire hotspot, each of the top 11, except for Hotspot4b_l77_s720724, covers at least part of its targeted domain and either overlaps or shares an interface with one of its targeted hotspot residues.

**Table 6.**
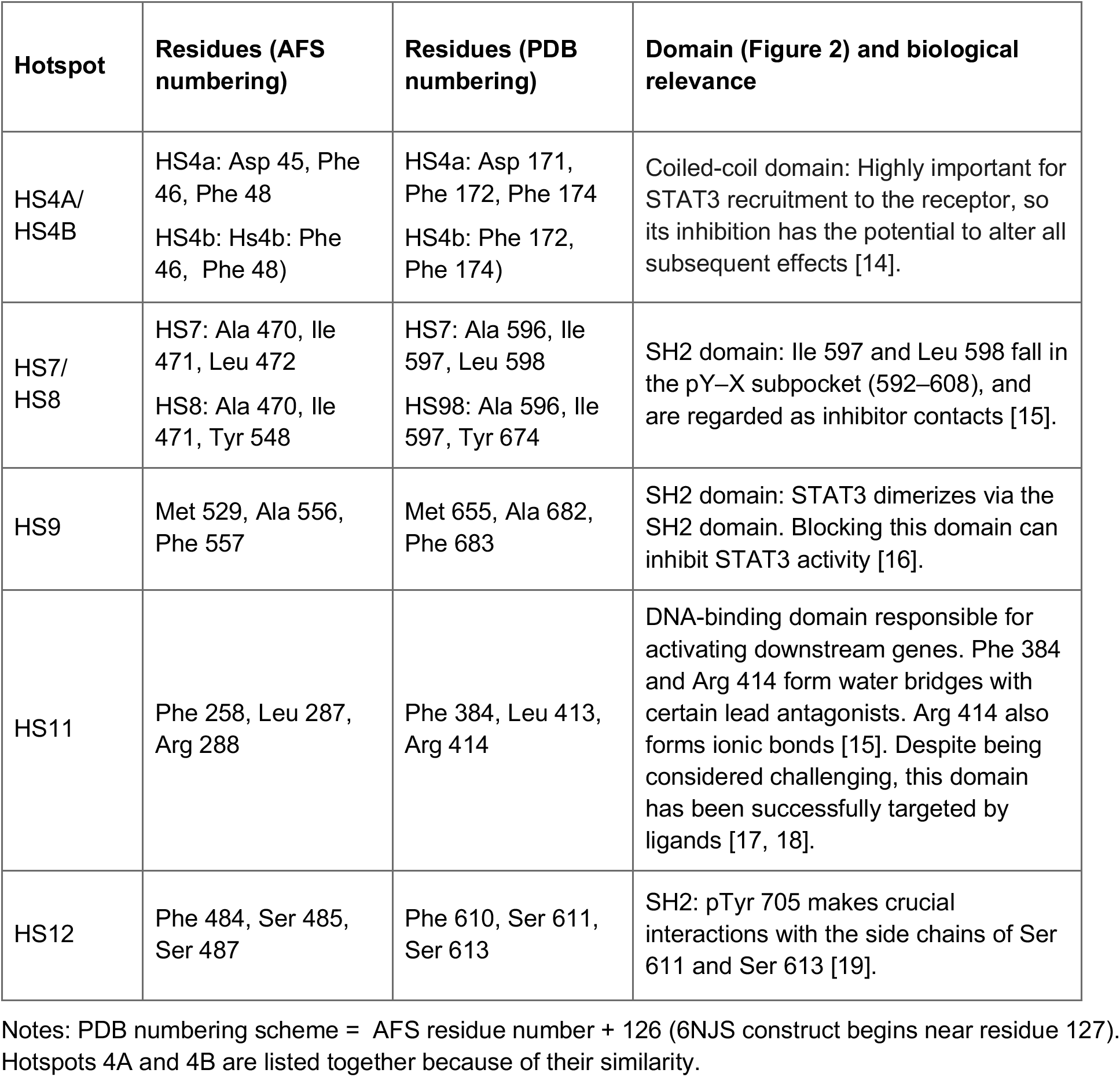
STAT3 domain and biological relevance of hotspots.

### 4.2 Targeted (hotspot) vs. simulated minibinder docking sites

Ten of the eleven binders dock within 4.5 Å of a hotspot residue, but visually, only four minibinders cover all their hotspot residues. The two measures of docking location often yield differing results. For example, the HS9 binder, 6NJS_HS9_l59_s417381, and two HS11 binders (6NJS_HS11_l64_s658467 and 6NJS_HS11_l53_s331773 l64, l53) covered all three of their hotspot residues visually, yet only one residue of each hotspot (M529 for HS9, R288 for HS11) was within 4.5 Å of the binder in question. The interface of the HS12 binder, 6NJS_HS12_l56_s197084, includes all three HS12 residues in the interface but visually, the binder doesn’t overlap those hotspot residues. Overlap without interface suggests a binder that sits over the hotspot without close contact. Interface without overlap suggests residues that are adjacent to the binder but not covered by it. Results also vary for a given hotspot. The two HS8 binders have 3/3 and 2/3 interface residues, and the five HS11 binders have 1–2 of 3.

HS12 was the most challenging hotspot. It produced only one passing binder from three campaigns. Its polar, charged pocket is likely a difficult target for BindCraft [20], compared to more hydrophobic patches.

### 4.3 Limitations

Several limitations apply to this study. First, no experimental validation was performed, and AF3 ipTM is a confidence score that doesn’t translate directly to a binding constant or other measure of biochemical affinity. We modeled the monomeric core fragment of STAT3 (6NJS), so surfaces involved in dimerization or DNA binding were modeled without their partners. Finally, we did not investigate the binding specificity of our minibinders against other proteins. Though it would be impossible to do so against all possible alternative targets, it would be instructive to evaluate the interaction of our minibinders with other STAT proteins, which share structural similarity with STAT3.

## 5. Conclusion

We used BindCraft to design minibinder proteins against seven surface hotspots on the coiled-coil, SH2, linker and DNA-binding domains of STAT3. From 139 designs that passed BindCraft’s filters, 98 had an AF2 ipTM ≥ 0.60 and were advanced to screening with AlphaFold3. From that, 11 designs generated against five hotspots met our high-confidence criterion of AF3 ipTM ≥ 0.75. All 11 binders are predicted to cover their targeted STAT3 domain. BLASTp found no match for six of the minibinders and only very low-significance matches for the other five, so the sequences appear to be absolutely novel.

We hope this set of minibinder protein candidates will help in evaluating AI-driven binder design for difficult targets such as STAT3.

## Author Contributions

E.R.S.:, data generation, data curation and analysis, visualization and figure generation, composing original draft, revision.

A.V.T.: conceptualization, supervision, instruction, methodology, revision and polishing.

## Acknowledgments

The authors thank Martin Pacesa for creating and posting the BindCraft pipeline for free use in Google Colaboratory.

## References

1. Samad, M.A., Ahmad, I., Hasan, A., Alhashmi, M.H., Ayub, A., Al-Abbasi, F.A., Kumer, A., Tabrez, S. STAT3 signaling pathway in health and disease. MedComm 6, e70152 (2025).

2. El-Tanani, M., Al Khatib, A.O., Aladwan, S.M., Abuelhana, A., McCarron, P.A., Tambuwala, M.M. Importance of STAT3 signalling in cancer, metastasis and therapeutic interventions. Cell. Signal. 92, 110275 (2022).

3. Tolomeo, M., Cascio, A. The multifaceted role of STAT3 in cancer and its implication for anticancer therapy. Int. J. Mol. Sci. 22, 603 (2021).

4. Yang, S., Wang, L., Chuan, Z., Nie, J. STAT3 inhibitor in oncology: a trial-based analysis of translational challenges and evolving therapeutic paradigms. Front. Oncol. 16, 1802640 (2026).

5. Jumper, J., Evans, R., Pritzel, A. et al. Highly Accurate Protein Structure Prediction with AlphaFold. Nature 596, 583–589 (2021).

6. Pacesa, M., Nickel, L., Schellhaas, C. et al. One-shot design of functional protein binders with BindCraft. Nature 646, 483–492 (2025).

7. Watson, J.L., Juergens, D., Bennett, N.R., Trippe, B.L., Yim, J., Eisenach, H.E., Ahern, W., Borst, A.J., Ragotte, R.J., Milles, L.F., Wicky, B.I.M., Hanikel, N., Pellock, S.J., Courbet, A., Sheffler, W., Wang, J., Venkatesh, P., Sappington, I., Vázquez Torres, S., Lauko, A., De Bortoli, V., Mathieu, E., Ovchinnikov, S., Barzilay, R., Jaakkola, T.S., DiMaio, F., Baek, M., Baker, D. De novo design of protein structure and function with RFdiffusion. Nature 620, 1089–1100 (2023).

8. Abramson, J., Adler, J., Dunger, J. et al. Accurate Structure Prediction of Biomolecular Interactions with AlphaFold3. Nature 630, 493–500 (2024).

9. Desai, D., Kantliwala, S.V., Vybhavi, J., Ravi, R., Patel, H., Patel, J. Review of AlphaFold 3: transformative advances in drug design and therapeutics. Cureus 16(7), e63646 (2024).

10. Gao, M., Skolnick, J. iAlign: a method for the structural comparison of protein-protein interfaces. Bioinformatics 26, 2259–2265 (2010).

11. Janin, J., Bahadur, R.P., Chakrabarti, P. Protein-protein interaction and quaternary structure. Q. Rev. Biophys. 41, 133–180 (2008).

12. Altschul, S.F., Madden, T.L., Schäffer, A.A., Zhang, J., Zhang, Z., Miller, W., Lipman, D.J. Gapped BLAST and PSI-BLAST: a new generation of protein database search programs. Nucleic Acids Res. 25, 3389–3402 (1997).

13. Choudhuri, S. Sequence alignment and similarity searching in genomic databases: BLAST and FASTA. In Bioinformatics for Beginners 133–155 (Academic Press, 2014).

14. Zhang, T., Kee, W.H., Seow, K.T., Fung, W., Cao, X. The coiled-coil domain of Stat3 is essential for its SH2 domain-mediated receptor binding and subsequent activation induced by epidermal growth factor and interleukin-6. Mol. Cell. Biol. 20, 7132–7139 (2000).

15. Mahalapbutr, P., Wonganan, P., Chavasiri, W., Rungrotmongkol, T. Butoxy mansonone G inhibits STAT3 and Akt signaling pathways in non-small cell lung cancers: combined experimental and theoretical investigations. Cancers 11, 437 (2019).

16. Cerulli, R.A., Shehaj, L., Tosic, I., Jiang, K., Wang, J., Frank, D.A., Kritzer, J.A. Cytosolic delivery of peptidic STAT3 SH2 domain inhibitors. Bioorg. Med. Chem. 28, 115542 (2020).

17. Anjum, S.M., Kumar, K.S., Umamaheswari, A., Lakhanpal, D., Swargam, S., Riazunnisa, K., Chandrasekhar, T. Identification of a natural antagonist for signal transducer and activator of transcription 3 (STAT3) by computational approach. Results Chem. 5, 100970 (2023).

18. Sabanés Zariquiey, F., da Souza, J.V., Estrada-Tejedor, R., Bronowska, A.K. If you cannot win them, join them: understanding new ways to target STAT3 by small molecules. ACS Omega 4, 13913–13921 (2019).

19. Fletcher, S., Turkson, J., Gunning, P.T. Molecular approaches towards the inhibition of the signal transducer and activator of transcription 3 (Stat3) protein. ChemMedChem 3, 1159–1168 (2008).

20. Pacesa, M. De novo binder design with BindCraft. BindCraft GitHub Wiki (2025). Accessed 22 Oct 2026.

21. Zouein, F.A., Altara, R., Chen, Q., Lesnefsky, E.J., Kurdi, M., Booz, G.W. Pivotal importance of STAT3 in protecting the heart from acute and chronic stress: new advancement and unresolved issues. Front. Cardiovasc. Med. 2, 36 (2015).

